# IL-10 Promotes Endothelial Progenitor Cell Driven Wound Neovascularization and Enhances Healing via STAT3

**DOI:** 10.1101/760165

**Authors:** Swathi Balaji, Emily Steen, Xinyi Wang, Hima V. Vangapandu, Natalie Templeman, Alexander J. Blum, Chad M. Moles, Hui Li, Daria A. Narmoneva, Timothy M. Crombleholme, Manish J. Butte, Paul L. Bollyky, Sundeep G. Keswani

**Affiliations:** Division of Pediatric Surgery, Department of Surgery, Texas Children’s Hospital and Baylor College of Medicine, Houston, TX, 77030, USA; Biomedical Engineering, Department of Biomedical, Chemical and Environmental Engineering, College of Engineering and Applied Sciences, University of Cincinnati, Cincinnati, OH, 45221, USA; Center for Children’s Surgery, Children’s Hospital Colorado and the University of Colorado School of Medicine, Aurora, CO, 80045, USA; Fetal Care Center Dallas, TX, 75230, USA; Division of Immunology, Allergy, and Rheumatology, Departments of Pediatrics and Microbiology, Immunology, and Molecular Genetics; University of California Los Angeles, Los Angeles, CA, 90095 USA; Division of Infectious Diseases, Department of Medicine, Stanford University School of Medicine, Stanford, CA, 94305, USA

**Keywords:** Wound healing, diabetes, IL-10, angiogenesis, endothelial progenitor cells, VEGF

## Abstract

Evidence from prior studies of cutaneous trauma, burns, and chronic diabetic wound repair demonstrates that endothelial progenitor cells (EPCs) contribute to *de novo* angiogenesis, anti-inflammatory reactions, tissue regeneration, and remodeling. We have shown that IL-10, a potent anti-inflammatory cytokine, promotes regenerative tissue repair in an adult model of dermal scar formation via the regulation of fibroblast-specific hyaluronan synthesis in a STAT3 dependent manner. While it is known that IL-10 drives EPC recruitment and neovascularization after myocardial infarction, its specific mode of action, particularly in dermal wound healing and neovascularization in both control and diabetic wounds remains to be defined. Here we show that IL-10 promotes EPC recruitment into the dermal wound microenvironment to facilitate neovascularization and wound healing of control and diabetic (db/db) wounds via vascular endothelial growth factor (VEGF) and stromal-cell derived factor 1 (SDF-1α) signaling. Inducible skin-specific STAT3 knockout (KO) mice were studied to determine whether the impact of IL-10 on the neovascularization and wound healing is STAT3 dependent. We found that IL-10 treatment significantly promotes dermal wound healing with enhanced wound closure, robust granulation tissue formation and neovascularization. This was associated with elevated wound EPC counts as well as increased VEGF and high SDF-1α levels in control mice, an effect that was abrogated in STAT3 KO transgenic mice. These findings were supported *in vitro*, wherein IL-10-enhanced VEGF and SDF-1α synthesis in primary murine dermal fibroblasts. IL-10-conditioned fibroblast media was shown to promote sprouting and network formation in aortic ring assays. We conclude that overexpression of IL-10 in the wound-specific milieu recruits EPCs and promote neovascularization, which occurs in a STAT3-dependent manner via regulation of VEGF and SDF-1α levels. Collectively, our studies demonstrate that IL-10 increases EPC recruitment leading to enhanced neovascularization and healing of dermal wounds.

## Introduction

Chronic ischemic wounds are prevalent and particularly debilitating in diabetic patients, as the impaired neovascularization seen in these wounds ultimately leads to recurrent infections, a failure to heal, and even extremity amputations, all of which impose major economic healthcare burdens.^1–3^ In contrast, physiologic dermal wound healing is characterized by a well-coordinated series of responses to injury, including granulation tissue formation, epithelial gap closure, inflammatory regulation, and robust neovascularization - a process initiated by local endothelial cells and by both circulating and tissue resident endothelial progenitor cells (EPCs) - to effectively restore circulation to the affected area and support tissue repair.^4^

Endothelial progenitor cells (EPCs) significantly contribute to wound neovascularization^5–7^ by differentiating into mature endothelial cells^8,9^ and forming vascular structures *de novo,* and via paracrine effects in the production of growth factors that support vascularization^10,11^. While the EPC phenotype continues to evolve,^12^ CD34, CD31/PECAM1, VEGFR2/Flk1/KDR, and CD133 are archetype cell markers to define these progenitor cells.^13,14^ More importantly, and consistent with the relevance of EPC- driven neovascularization in tissue repair, impaired mobilization and recruitment of EPCs has been implicated in diabetic wounds.^15–25^ Therefore, it is important to develop innovative strategies aimed at improving molecular signals that trigger EPC mobilization, homing, survival, and function as a means to ultimately achieve a meaningful clinical impact in the treatment of chronic wounds.^26–30^

One of the factors that mobilize EPCs from the bone marrow (BM) into circulation after injury is vascular endothelial growth factor (VEGF),^31–33^ produced in injured tissues in response to hypoxia.^34^ VEGF is known to activate BM matrix metalloproteinase-9 (MMP-9), which in turn releases stem cell factor (SCF) to facilitate EPC proliferation and mobilization from the BM niche into the peripheral circulation.^35^ Furthermore, recent studies have shown that VEGF promotes stromal cell-derived factor 1 alpha (SDF-1α) and CXC chemokine receptor 4 (CXCR4)-dependent signaling, leading to homing of bone marrow hematopoietic progenitor and stem cells and cardiac stem cells to tissue repair sites in models of myocardial infarction.^36,37^ This signaling pathway is also known to be involved in EPC homing to sites of hypoxia/ischemia.^38,39, 40–42^ Previous studies demonstrated that fibroblasts are likely a critical source of VEGF and SDF-1α production in cutaneous wounds.^43,44^ Existing evidence indicates that SDF-1α is constitutively expressed by BM stromal cells and in so doing plays a key role in maintaining the BM niche, as well as providing directional cues that orchestrate CD34+ cell homing. Under homeostatic conditions, high SDF-1α expression in the BM is considered a strong chemotactic factor whose function is to retain CD34+ progenitor cells within the BM niche. After an injury, SDF-1α is released by the injured tissue, ^44,45^ and the higher local levels of SDF-1α stimulates mobilization of progenitor cells out from the bone marrow and their recruitment to the injury site. Hattori et al. demonstrated that elevated SDF-1α levels in peripheral blood resulted in the mobilization of these cells to the peripheral circulation.^42^ In support of this finding, it has been shown that EPCs can be recruited from the bone marrow into peripheral hypoxic/ischemic tissue sites in response to treatment with SDF- 1 α and/or VEGF.^46^ However, the high concentrations and multiple doses needed for individual angiogenic growth factors to effect a relevant clinical outcome are problematic. In addition, the growth and chemotactic factors such as VEGF, granulocyte colony stimulating factor (G-CSF), SCF, and SDF-1α that have been used for EPC mobilization have also been shown to induce hematopoietic stem and progenitor cell, as well as monocyte and macrophage, mobilization; by increasing the inflammatory burden at the tissue repair site, this undue mobilization may have negative effects on wound repair and healing.^47^ In other words, while it is plausible that angiogenic growth factor treatment improves progenitor cell homing and EPC-mediated vascular remodeling, it is equally conceivable that it may affect the survival and/or function of EPCs at the sites of tissue repair due to increased inflammation.

Notably, a recent study of a murine myocardial infarction model showed that IL-10, a potent anti-inflammatory cytokine,^48,49^ facilitates EPC mobilization and revascularization.^45^ Moreover, a study in which IL-10-transfected EPCs were adoptively transferred into the retinal microenvironment of diabetic rats showed improved vascular repair.^50^ Although these collective findings support the role of IL-10 in EPC-mediated neovascularization, the mechanisms that determine such a function in cutaneous wounds remain to be elucidated. We and others have shown that, in non-diabetic wound healing, IL-10 overexpression in postnatal cutaneous dermal wounds induces regenerative (scarless) tissue repair via inflammatory regulation^51–53^ and the fibroblast-mediated formation of a extracellular wound matrix rich in hyaluronan^54^ through a STAT3- dependent signaling pathway.^55,56^. The ability of IL-10 to drive EPC recruitment and neovascularization, along with its potential to promote dermal wound closure, have been largely unstudied.

We hypothesized that, upon injury to the skin, IL-10 stimulates the local expression of VEGF and SDF-1α to mobilize and recruit EPCs into cutaneous wounds in a STAT3-dependent manner, inducing neovascularization and promoting wound healing. To obtain experimental evidence that supports our hypothesis, we studied IL-10 overexpression in non-diabetic and diabetic murine models of dermal wound healing using a previously described lentivirus (LV) transduction approach.^51,54^ To assess the proposed contribution of STAT3 signaling, we investigated IL-10 mediated tissue repair using a recently developed skin-specific STAT3 knockdown mouse model.^54^

## Materials and Methods

### Generation of inducible STAT-3 knockout transgenic murine model

All animal procedures were approved by Cincinnati Children’s Hospital Medical Center and Baylor College of Medicine Institutional Animal Care and Use Committee. 8- 12 week old wild type (WT) mice (C57BL/6J; hereafter referenced as WT), type II diabetic mice (BKS.Cg-m+/+Lepr^db^/J; hereafter referenced as db/db), MMP-9 knockout (KO) mice (FVB.Cg-Mmp9tm1Tvu/J) and strain-matched WT mice (FVB/NJ) were purchased from Jackson laboratories (Bar Harbor, ME). Db/db mice with serum glucose >400 mg/dl and weight >40gms were used in all experiments. Conditional STAT3 KO mice were developed as explained previously.^54^ Briefly, B6.Cg-Tg(UBC-cre/ERT2)1Ejb/J Cre-expressing mice driven by the human ubiquitin C (UBC) promoter were bred to mice containing a loxP-flanked STAT3^flox/flox^ sequence (Jackson Laboratories). Double transgenic phenotype (STAT3^Δ/Δ^) was confirmed by genotyping^54^ and 8-12 week old mice were used in all experiments. The dorsal skin was shaved and 4-OHT was topically administered (10 mg/ml in sterile vegetable oil, 150 µl every day for seven days) to activate Cre-mediated recombination and achieve STAT3 deletion (STAT3^-/-^) in the dorsal skin, with vegetable oil administration alone serving as vehicle control group (STAT3^Δ/Δ ctrl^). STAT3 knockdown in the skin was quantified using Western blotting and band densitometry on dorsal skin snips from treated area as described previously, ^54^ prior to using the mice in excisional wounding.

### Excisional wounding and tissue harvest

Mice were anesthetized with isoflurane inhalation (0.5 ml, titrated). Dorsal skin was shaved and prepared by scrubbing alternately with isopropyl alcohol and povidone-iodine. Mice were divided into two treatment groups, where the left side dorsal flank was pre-treated with either 50µl of 1X10^6^ TU/ml of lentiviral IL-10/GFP or lentiviral GFP alone, injected intradermally and labeled with India ink (n=4-6 mice/treatment group at each time point; both male and female). PBS labeled with India ink was injected in the bilateral flank as an internal control. A third group of mice, where both the bilateral flanks were pre-treated with intradermal injection of PBS labeled with India ink, served as the external control treatment group. After 4 days to allow transgene expression, mice were anesthetized and two full thickness excisional wounds extending to the panniculus carnosus were created on both flanks at the treatment sites using a 4mm biopsy punch (Miltex, Plainsboro, NJ). To prevent skin contraction, 6mm silicone stents were secured around the wounds with skin glue and 6-8 interrupted sutures with 6-0 nylon (Ethicon Inc., Somerville, NJ). Stented wounds were covered with Tegaderm^TM^ (3M, St. Paul, MN). In STAT3 mice cohorts, daily topical administration of 4-OHT or vegetable oil was continued. Specifically in MMP9^-/-^ mice, in order to rescue the deficiency in release of SCF caused by lack of MMP9 and to reinstate endothelial progenitor cell mobilization, 0.5 ug/kg of SCF (Peprotech, Rocky Hill, NJ) was injected daily post wounding via tail vein.^47^

At day three and day seven post-wounding, mice were euthanized and peripheral blood, wounded skin, and femur and iliac bones were collected. The bone marrow was flushed out with saline at stored at −80°C. Wounds were bisected, and one half was fixed in 10% formalin, and paraffin-embedded. The other half was processed for RNA and protein isolations.

### Histologic evaluation of wound morphology

5 µm wound sections were cut from paraffin blocks. Epithelial gap closure and granulation tissue deposition were analyzed via Hematoxylin and Eosin (H&E) staining and morphometric image analysis using Nikon Elements (Nikon Instruments, Melville, NY). Wound ECM composition was determined using Movat’s Pentachrome staining as per the manufacturer’s protocol (Polyscientific, Bay Shore, NY).

### Immunohistochemical Staining

5 µm serial sections were dewaxed in three changes of xylene for 10 minutes each and rehydrated in an ethanol to distilled water series. Antigen retrieval was performed using 1X Target Retrieval Solution (Dako, Carpinteria, CA) at 95°C for 20 minutes followed by a cool down to room temperature. Following a five minute wash in distilled water, sections were permeablized in 1X Phosphate Buffered Saline with 0.1 % Tween (PBSTw) for 10 minutes. Endogenous peroxidase was blocked with 3% H2O2 followed by blocking of nonspecific protein binding with a solution of 5% Rabbit Serum + 1% Bovine Serum Albumin in PBSTw for 2 hours at room temperature. A M.O.M kit (Vector Laboratories, Burlingame, CA) was applied to slides per manufacturer’s instructions if mouse monoclonal primary antibodies were used. Immunostaining with antibodies against CD45 (1:2500, Abcam ab10558, Cambridge, MA), Lamp2a (1:50; Abcam ab18528), MECA-32 (1:10, deposited to the Developmental Studies Hybridoma Bank, University of Iowa by Butcher), STAT3 (1:500, Cell Signaling Technology 9139, Danvers, MA), Phospho-STAT3 (Tyr705) (1:200, Cell Signaling 9145), TGF-β1 (1:100, Abcam ab92486), and TGF-β3 (1:100, Abcam 15537) was performed for one hour at room temperature, followed by incubation with biotinylated species specific secondary antibodies (1:200, Vector Laboratories). ABC-DAB peroxidase based staining followed by hematoxylin counter staining (Vector Laboratories) was performed, and slides were dehydrated and mounted in xylene based permanent mounting media.

Endothelial progenitor cells (EPCs) were identified using double immunofluorescent staining for CD133 and Flk-1. 5 µm paraffin sections were prepared similar to above and autofluorescence was blocked using 50mM Ammonium Chloride for 20 minutes. Following a rinse in PBSTw, nonspecific protein binding was blocked with a solution of 5% Normal Goat Serum, 1% Bovine Serum Albumin in PBSTw for two hours at room temperature. Primary antibodies were labeled according to the Zenon Complex Formation protocol (Invitrogen, Carlsbad, CA). A 3:1 molar ratio of Fab to antibody target was used for CD133 (AbCam ab19898), and 6:1 molar ratio for FLK-1 (AbCam, ab2349). The CD133-488 complex was diluted 1:250 while the FLK-1-647 complex was diluted 1:25 in block and incubated for one hour at room temperature. The sections were then washed three times in PBSTw for 5 minutes. Following the washes, the anibody-Fab complex was fixed to the samples using 4% paraformaldhyde for 15 minutes. The samples were then washed three times in PBS, and mounted using Prolong Gold plus DAPI (Invitrogen).

The average number of CD45+, Lamp2a+ and CD133+Flk-1+ cells was quantified by counting positive cells in six high power fields (HPF, 40X) per wound section. Capillary lumen density was measured as the average number of MECA32-positive lumens from six HPF (40X) per section. HPF were equally distributed between the two wound edges.

### Flow cytometry

350 µl of whole blood per mouse was collected. Red blood cells were lysed using lysis solution per manufacturer’s instructions (Qiagen, Alameda, CA). The cells were washed and centrifuged twice in 10ml of 1% BSA-PBS and re-suspended in 1ml of 1% BSA-PBS solution and counted. EPCs were co-labeled using APC-conjugated anti-CD34 (1µl per 5×10^6^ cells), FITC-conjugated anti-CD133 (1.5µl per 1X10^6^ cells), and PE-conjugated anti-Flk-1 (2µl per 1×10^6^ cells) monoclonal antibodies (BD Biosciences, San Jose, CA) in the dark for 20 minutes at 4°C with gentle rocking. Unstained control and individual color controls were also included for gating. Cells were washed and re- suspended in 350µl of medium containing 7-AAD viability stain for live/dead discrimination (3µl per 1×10^6^ cells, eBioscience, San Diego, CA). Using the FACS Canto-II flow cytometer (BD Biosciences, San Jose, CA), single cells were gated to obtain a live CD34- positive population. Within this population, cells that co-expressed CD133 and Flk-1 were counted as EPCs. Each data point included at least 1000000 events. Flow data were then analyzed using FlowJo software (Tree Star Inc., Ashland, OR) by a blinded investigator.

### Primary Cell culture experiments

Primary adult (8-10 week old) murine dermal fibroblasts were isolated from the skin of C57BL/6J mice (Jackson Laboratories, Bar Harbor, ME, USA) using standard isolation protocols.^57^ The fibroblast culture was routinely maintained at 37°C under 5% CO_2_ in a humidified chamber and cultured in Dulbecco’s Modified Eagle’s media (DMEM; GIBCO, Carlsbad, CA, USA) supplemented with 10% bovine growth serum (BGS; Hyclone, Logan, UT, USA), 100U penicillin, 100µg streptomycin + 0.25µg amphotericin B (PSF; Invitrogen, Carlsbad, CA, USA). All cells used were between passages 5-10.

Fibroblasts were seeded at 2×10^5^ cells/well in a six-well plate in DMEM containing 10% BGS and allowed to settle overnight. Cells were then serum starved in DMEM culture media with 2% BGS and the conditioned culture was supplemented with 200ng/ml of murine IL-10 recombinant protein (Peprotech, Rocky Hill, NJ). After 48 hours, supernatant was collected to investigate the effect of IL-10 stimulation on VEGF and SDF-1α production by the fibroblasts. Cells from two different passages were tested in triplicate experiments.

### Aortic Ring Assay

An aortic ring assay was performed as described previously.^58^ Briefly, thoracic aortas were dissected from 8-10 week old male and female C57BL/6J mice after euthanasia. The periaortic connective tissue was carefully removed under a dissecting scope (M80 Stereomicroscope, Leica Microsystems, Buffalo Grove, IL) without damaging the vessel wall. The aorta was then cut into 0.5mm wide rings using a #10 blade and affixed on a culture dish with a thin layer of growth factor-reduced basement membrane matrix Matrigel (Corning, NY) and covered with DMEM culture media. These rings were first serum-starved overnight in DMEM containing no serum to equilibrate their growth factor responses. Aortas were grown in (1) DMEM complete media similar to the primary culture described earlier, (2) DMEM complete media supplemented with 200ng/ml of murine IL-10 recombinant protein, (3) conditioned media from untreated adult dermal fibroblasts, or (4) conditioned media from adult dermal fibroblasts treated with 200ng/ml of murine IL-10 recombinant protein. Conditioned media for conditions 3 and 4 was from primary cell culture experiments used for detection of growth factor production as described above. Three rings per treatment were studied and the experiment was repeated two times with aortas from different mice and conditioned media from different primary cell isolations. The media was changed every 2-3 days. Sprout outgrowth was monitored on alternate days using phase contrast imaging (Leica DMi8). Image analysis software LASX (Leica) was used to measure the relative sprouting area (calculated as the difference in ring area between day 1 and 12).

### Quantitative Real-Time PCR

Wound tissue was homogenized in RLT buffer and total RNA was isolated per manufacturer’s recommendations (mini kit, Qiagen, Valencia, CA, USA). cDNA was synthesized from 1µg of RNA using the High Capacity cDNA Reverse Transcription kit (Applied Biosystems, Foster City, CA, USA) following manufacturer’s protocols. SYBR green assays were designed to span intron/exon boundaries. Oligonucleotides were aligned against the mouse genome by Primer-BLAST (www.NCBI.org) to ensure specificity. Gene expression was assayed in triplicate using 1/40^th^ of the cDNA template, and 300nM of forward and reverse primer in a 25µl Power SYBR Green PCR Master Mix reaction in the StepOne-Plus Real-Time PCR System (Applied Biosystems). Gene expression was normalized to mouse RPS29 gene expression. Relative expression values were calculated using the Comparative Ct (ΔΔCt) method.^59^ Oligonucleotide primer sequences used were as follows:

VEGFa 1F 5’-TTAAACGAACGTACTTGCAGATG-3’ and 1R 5’- AGAGGTCTGGTTCCCGAA-3’

SDF-1α 1F 5’-GGTTCTTCGAGAGCCACATCG-3’ and 1R 5’- ACGGATGTCAGCCTTCCTCG-3’

Rps29 1F 5’-TCTGAAGGCAAGATGGGTCAC-3’ and 1R 5’- GTGGCGGTTGGAGCAGACG-3’.

### Enzyme Linked Immunosorbent Assays (ELISA)

Wound tissue was homogenized in 50mM Tris-HCl buffer containing 1% NP40, aprotinin (3.3μg/ml), leupeptin (10μg/ml) and pepstatin (4μg/ml) and stored at −80°C until testing (Sigma-Aldrich, St. Louis, MO). VEGF and SDF-1α protein levels were determined using Quantikine ELISA Kits (R&D Systems, Minneapolis, MN) per manufacturer’s instructions. ELISA data was normalized to total protein in each sample, calculated using the Coomassie Plus protein assay (Thermo Scientific, Logan, UT).

### Statistical analyses

Data was analyzed using ANOVA followed by Tukey post-hoc means comparison test. The data are expressed as mean±standard deviation. Differences at p<0.05 were considered to denote statistical significance.

## Results

### IL-10 overexpression improved wound healing and reduced inflammatory cell infiltration via a STAT3-dependent mechanism

We sought to determine the effect of IL-10 overexpression on the early wound healing responses, including wound closure, re-epithelialization, granulation tissue formation, and neovascularization, which have not been previously studied. To further determine whether these effects of IL-10 were mediated via conserved STAT3 signaling, we used a tamoxifen (4-OHT)-inducible cre driven skin-specific STAT3 knockout murine model. We generate the STAT3^Δ/Δ^ transgenic mice as described previously^54^. Topical application of 4-OHT in vegetable oil as a vehicle down regulated STAT3 levels in the skin of these mice, compared to topical application of vegetable oil vehicle alone, which served as vehicle control for all experimental purposes (Figure S1).

Prior to assessing the effect of IL-10 overexpression on wound healing, we first compared baseline wound healing with PBS (moist wound) control treatment in the three groups of mice, i.e. C57BL/6J (WT), STAT3^Δ/Δ Ctrl^, and STAT3^-/-^, and showed that at day three, epithelial gap closure and granulation tissue formation in STAT3^Δ/Δ ctrl^ transgenic mice was comparable to WT mice. Further, these parameters were not affected in STAT3^-/-^ mice wounds (Figure 1g, h; dotted line represents PBS wounds in WT). These data confirm that the ∼60% STAT3 knockdown we achieved at the protein levels in the skin of 4-OHT-treated transgenic mice has no impact on the baseline ability of these mice to heal cutaneous wounds.

**Figure 1:**
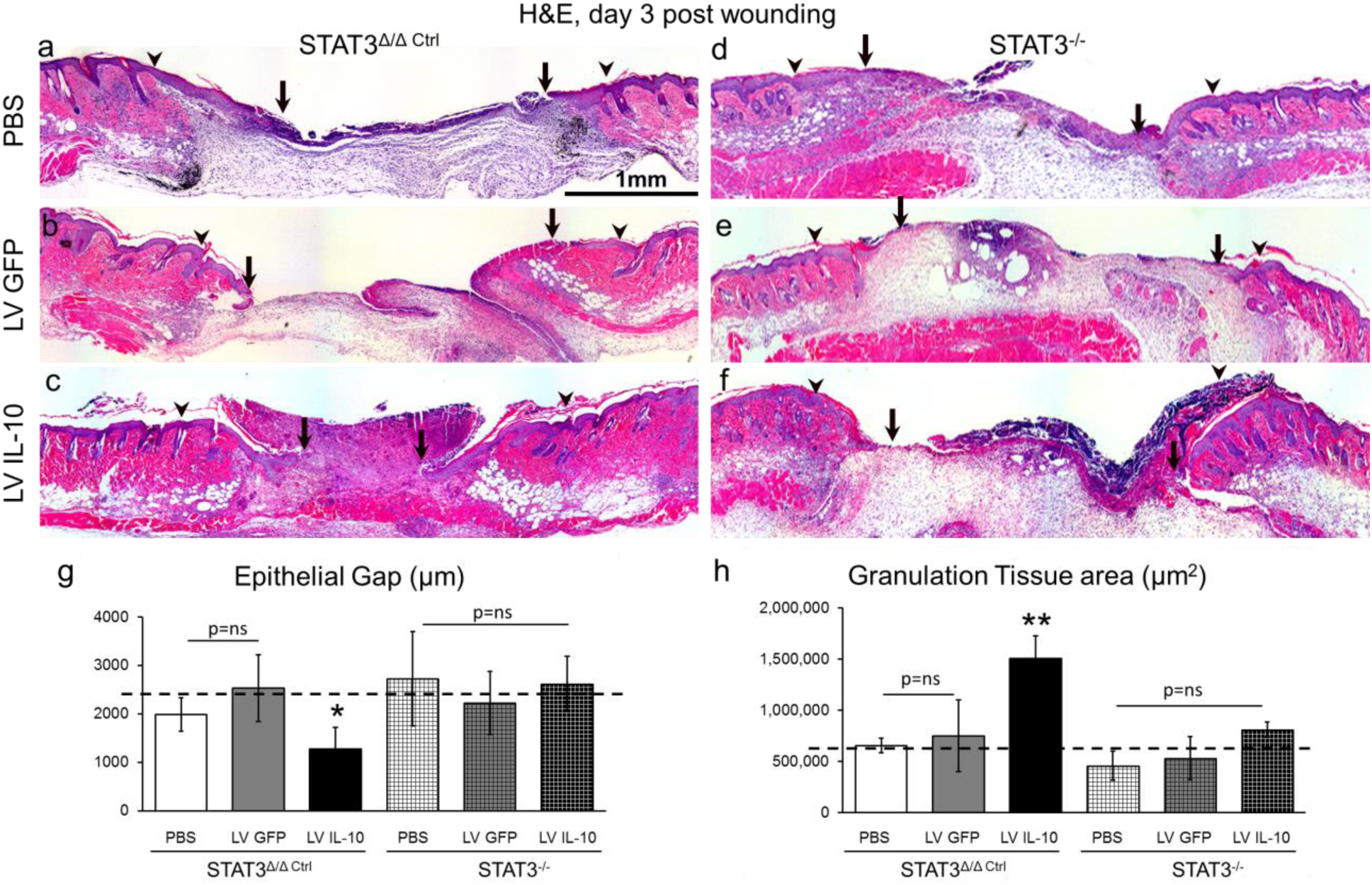
IL-10 overexpression enhanced wound closure in a STAT3 dependent manner *in vivo*. (a-f) Hematoxylin and Eosin staining of wounds at day 3 post wounding. Wounds are marked in India ink; wound margins are indicated by arrowheads and the encroaching epithelial margins are indicated by arrows. (a-f) Original magnification: 4X, Scale Bar=500µm; As compared to PBS (a) or lentiviral GFP (b) control treatments, lentiviral IL-10-treated wounds show enhanced wound closure and robust granulation tissue formation (c) in STAT3^Δ/Δ ctrl^ vehicle control mice. IL-10’s effects on wound healing were abrogated in STAT3^-/-^ mice, as evidenced by significantly impaired wound morphology (d-f). (g-h) Quantitation of epithelial gap (g) and granulation tissue (h) amongst the different treatments based on image analysis show that while there is no significant difference in gap closure between PBS and lentiviral GFP wounds, lentiviral IL-10 treated wounds exhibit increased gap closure and granulation tissue formation, whereas in STAT3^-/-^ mice this effect is abrogated. Bar plots: mean ± SD, 2 sections/ wound, n=4 wounds from different mice/ treatment group, *** p<0.001 by ANOVA.

Next, we evaluated the effect of lentiviral-transduced IL-10 (LV IL-10) overexpression on wound healing at day 3 post wounding. Similar to our previous reports, where we demonstrated efficient transduction with transgenic protein expression detected at the base of wounds within 72 h of treatment, we saw a significant increase in IL-10 levels at day 3 post-wounding as well as enhanced phosphorylated-STAT3 signaling in STAT3^Δ/Δ ctrl^ transgenic mice wounds treated with LV IL-10 (Figure S2). Histologic observation and morphometric analysis of day 3 wounds in STAT3^Δ/Δ ctrl^ mice demonstrated that IL-10 overexpression significantly enhanced the speed of epithelial gap closure and was associated with robust granulation tissue formation. LV GFP and PBS control wounds, in comparison, had a paucity in granulation tissue and reduced travel distance of the encroaching epithelial margins (Figure 1a-c; g-h, arrow heads indicate edge of the punch wound, arrows indicate the tip of the encroaching epithelial tongues). As expected, the effects of IL-10 were abrogated in STAT3^-/-^ mice (Fig. 1f), which had comparatively similar epithelial gap closure and granulation tissue formation to that observed in LV GFP or PBS wounds (Fig. 1d-h), providing support for the postulate that IL-10’s effects are mediated via downstream STAT3 signaling.

At day 7 the extent of epithelial gap closure was similar between PBS-, LV GFP-, and LV IL-10-treated STAT3^Δ/Δ ctrl^ mice (Figure 2a-c). However, at the same time point, IL-10-treated wounds demonstrated more organized cell layers in the epidermis, and robust granulation tissue (Figure 2g), with uniform distribution of the cellular density throughout the healing wound. Moreover, wounds treated with LV IL-10 had a greater degree of collagen deposition in the wound center (Figure S3). In contrast, LV GFP- (Figure 2b and e) and PBS-treated (Figure 2a and d) wounds demonstrated dense cellularity only at the encroaching edges, with lower cell density and collagen staining at the wound center, indicating a slower progress of wound healing. As anticipated, LV IL- 10 treatment also significantly decreased the CD45+ (Figure 2h) and Lamp2a+ (Figure 2i) inflammatory cell burden in the wound compared to LV GFP and PBS controls. Similarly, immunohistochemical evaluation of TGF-β1 and -β3 staining showed that LV IL-10-treated wounds had comparable TGF-β1 wound bed expression patterns to those of control wounds (Figure 2j), but higher TGF-β3 wound bed expression (Figure 2k). Conversely, the previously-seen effect of IL-10 on granulation tissue formation (Figure 2d-g), inflammatory cell infiltration (Figure 2h-i) and TGF-β3 expression (Figure 2k) was largely abrogated in STAT3^-/-^ mice and resulted in a wound healing response similar to LV GPF and PBS control wounds. Taken together, these data support our prediction that IL-10 overexpression improves initial wound closure and granulation tissue deposition by mechanisms that are STAT3 dependent, ultimately leading to superior remodeling and regeneration phenotype that we have recently reported.^54^

**Figure 2:**
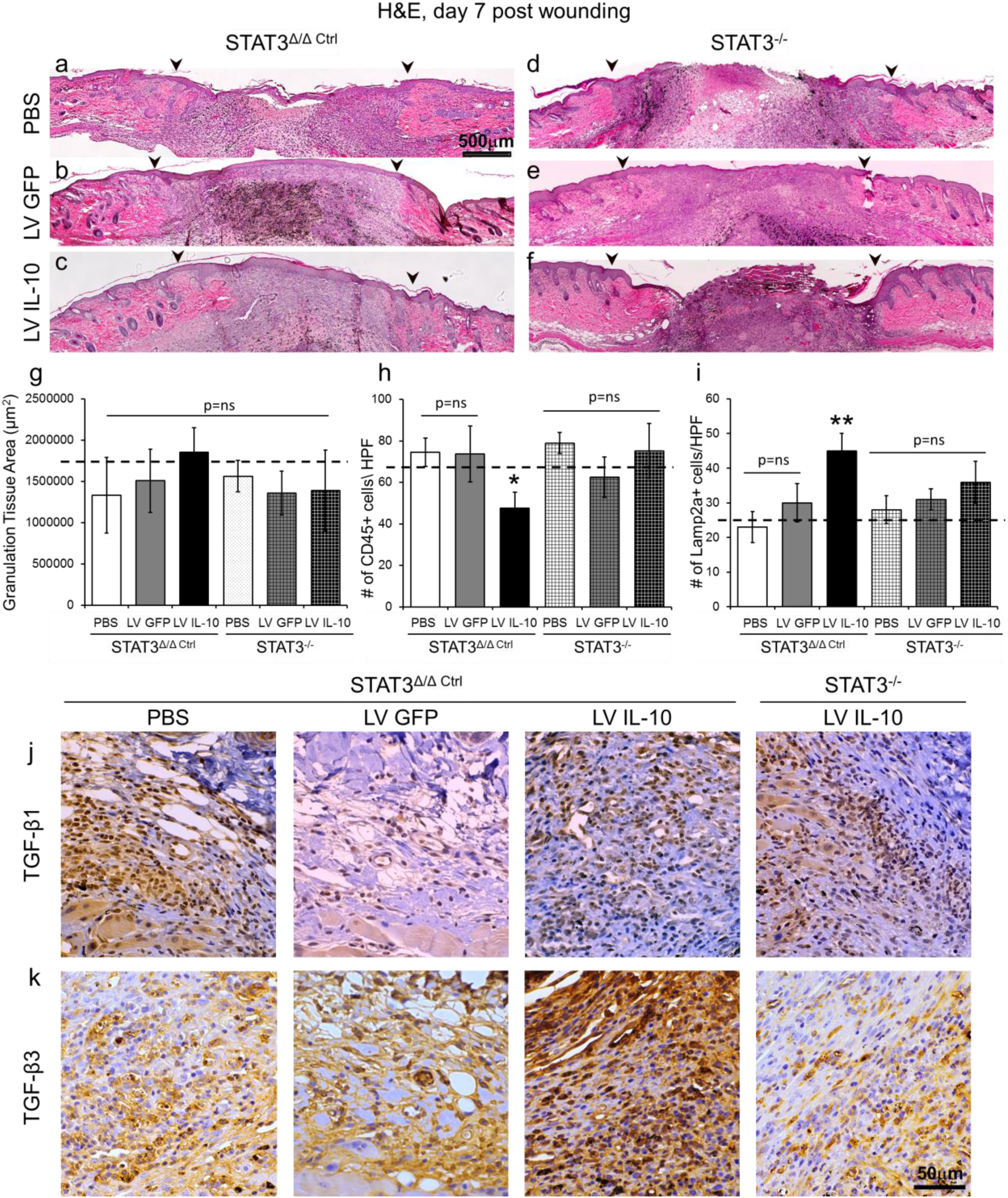
IL-10 overexpression reduced inflammation and enhanced wound remodeling and regenerative phenotype in a STAT3 dependent manner. (a-f) Hematoxylin and Eosin staining of wounds at day 7 post wounding. Wounds are marked in India ink; wound margins are indicated by arrowheads. (a-f) Original magnification: 4X, Scale Bar=500µm; (a-c) similar gap closure is observed between the three groups – PBS, lentiviral GFP and lentiviral IL-10 treated wounds in STAT3^Δ/Δ ctrl^ mice. However, IL-10 treated cohort display thicker epidermis and robust granulation tissue as compared to the control treatments.(d-f) All treatment cohorts healed similarly in STAT3^-/-^ mice and the effect of IL-10 on the epidermis and granulation tissue deposition is not apparent. (f) quantitation of granulation tissue between different treatment groups in STAT3^Δ/Δ ctrl^ show a modest trend in IL-10 treated mice, however, with no statistical significance between STAT3^Δ/Δ ctrl^ and STAT3^-/-^ with different treatments. (h) quantification of staining with CD45 show significantly lower CD45+ cells per high power field (HPF; 40X) in lentiviral IL-10 wounds as compared to lentiviral GFP or PBS treatments in STAT3^Δ/Δ ctrl^ mice, which is abrogated in STAT3^-/-^ mice. (i) Elevated expression of Lamp2a positive cells was observed in IL-10 treated wounds as compared to the control treatments in STAT3^Δ/Δ ctrl^ mice, which is abrogated in STAT3^-/-^ mice. (j) Panels left to right show similar TGF-β1 staining pattern in controls vs IL-10 treated wounds in STAT3^Δ/Δ ctrl,^ mice, that remain similar in expression in STAT3^-/-^ wounds. (k) A definite increase in TGF-β3 expression is seen in IL-10 treated wounds as compared to control treated ones in STAT3^Δ/Δ ctrl^ mice, however, this effect induced by IL-10 waned in STAT3^-/-^ wounds. Scale bar=50µm in (j- k); Bar plots: mean ± SD, 2 sections/ wound, n=4 wounds from different mice/ treatment group, *** p<0.001 by ANOVA.

### IL-10 overexpression increased capillary density and EPC infiltration of the wound tissue via STAT3

We then investigated the effect of IL-10 overexpression on wound capillary density and EPC infiltration. To that end, we performed immunohistochemical staining with MECA-32, a panendothelial cell antigen: LV IL-10-treated wounds in STAT3^Δ/Δ ctrl^ mice developed vessels with well-defined lumens at day 7, in contrast to control LV GFP- or PBS-treated wounds, which showed mostly non-cohesive individual cell staining (Figure 3a-c). LV IL-10-treated wounds also showed significantly increased capillary lumen density per high power field (LV IL-10: 24.17±2.69 vessels/HPF vs. LV GFP: 15.7±2.49, PBS: 14.6±3.8, p<0.001; Figure 3m). In contrast, STAT3^-/-^ wounds treated with LV IL-10 failed to achieve the anticipated effects on capillary lumen density (STAT3^-/-^ + LV IL-10: 12.18±4.18 vessels/HPF vs. STAT3^Δ/Δ ctrl^ + LV IL-10: 24.17±2.69, p<0.001; Figure 3d-f, m).

**Figure 3:**
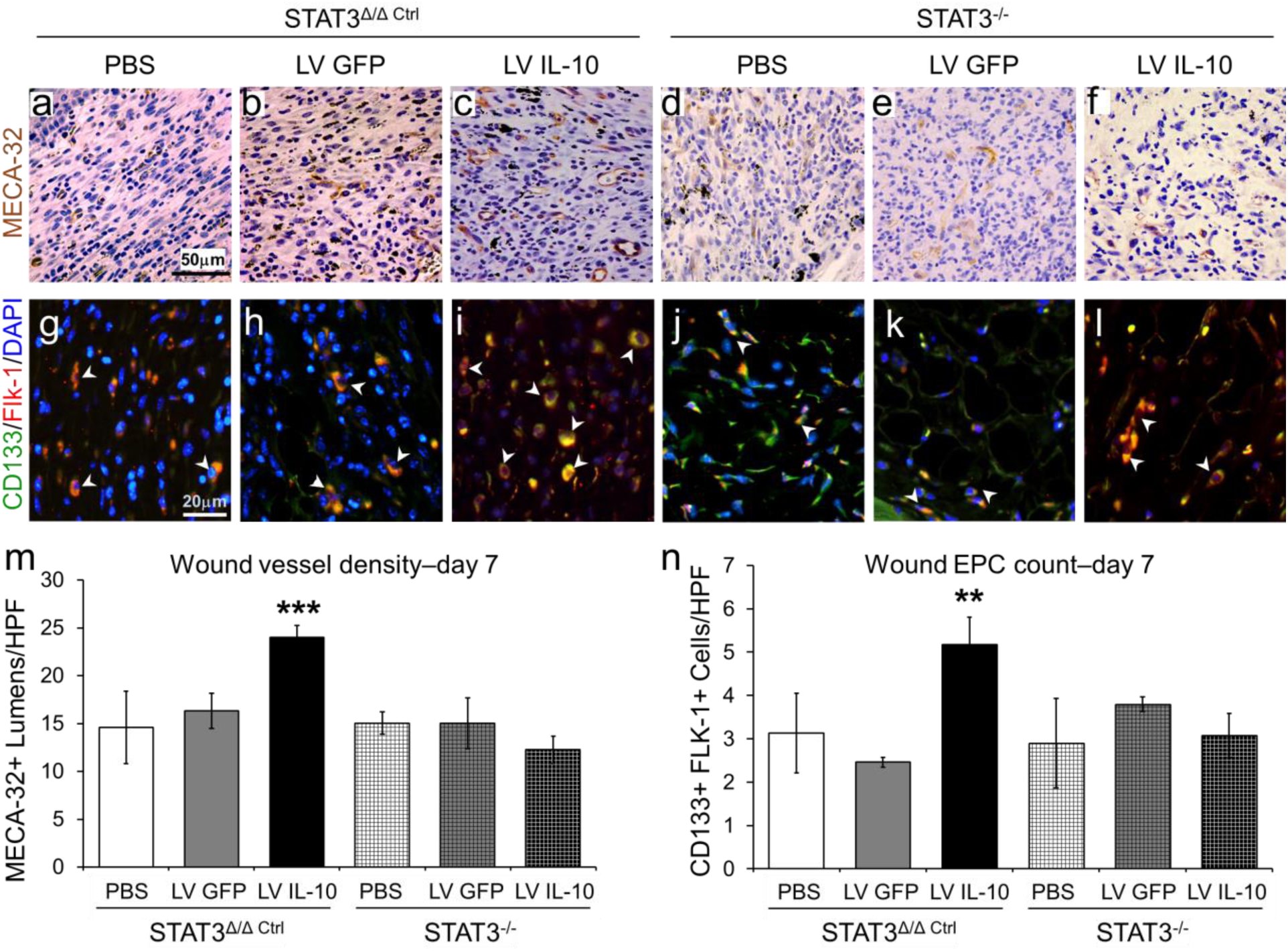
IL-10 overexpression enhanced wound neovascularization and EPC infiltration in a STAT3 dependent manner. (a-f) Capillary lumens are marked by MECA- 32 staining. (g-l) EPCs are identified by their characteristic morphology, large eccentric nuclei and CD133+/FLK-1+ staining (CD133–green, FLK-1–red, merged–yellow). Representative MECA-32+ and CD133+/Flk-1+ wound images for each treatment group at day 7 are shown here. Lentiviral IL-10 treatment significantly enhanced wound vessel density (m) and EPC numbers (n) per high power field (HPF), compared to lentiviral GFP or PBS treatments in STAT3^Δ/Δ ctrl^ mice, which was abrogated in STAT3^-/-^ mice. CD133+/FLK-1+ EPCs are indicated by arrow heads. Bar plots: mean±SD. 2 sections/wound, n=3-4 wounds from different mice/ treatment group, *** p<0.001 by ANOVA. Scale Bars=50µm and Original magnification: 40X in (a-f); Scale Bars=20µm and Original magnification: 80X in (g-l).

Similarly, LV IL-10-treated wounds in STAT3^Δ/Δ ctrl^ mice had significantly increased infiltration of CD133^+^/Flk-1^+^ EPCs, as shown by immunofluorescence co-labeling at day 7 after wounding (LV IL-10: 5.17±1.03EPCs/HPF vs. LV GFP: 2.43±0.97, PBS: 3.13±1.42, p<0.01; Figure 3g-i, n). In contrast, LV IL-10 once again failed to achieve the anticipated increase in wound EPC expression when administered to STAT3^-/-^ mice, wherein the EPC count was no different than that quantified in LV GFP- or PBS-treated control wounds (STAT3^-/-^ + LV IL-10: 3.38±1.63 EPCs/HPF vs. STAT3^Δ/Δ ctrl^ + LV IL-10: 5.17±1.03, p<0.01; Figure 3j-l, n). These data further support the significant role of IL-10 in driving both neovascularization and EPC recruitment to wounds as part of its function in regulating cutaneous wound healing, as well as underscoring the dependence of this process on STAT3 signaling.

### IL-10 overexpression increased EPC mobilization in response to wounding via a STAT3-dependent mechanism

We then determined whether the increase in EPC counts in LV IL-10-treated wounds was facilitated by an increase in EPC mobilization. As we have previously shown that that in response to cutaneous wounding, EPC levels peak in peripheral blood at day 3 post-injury^60^, we evaluated the peripheral blood for the presence of CD34^+^CD133^+^Flk- 1^+^ co-labelled cells in STAT3^Δ/Δ ctrl^ and STAT3^-/-^ mice after treatment of their cutaneous wound with LV IL-10, LV GFP and PBS using flow cytometry (Figure 4a-c).

**Figure 4:**
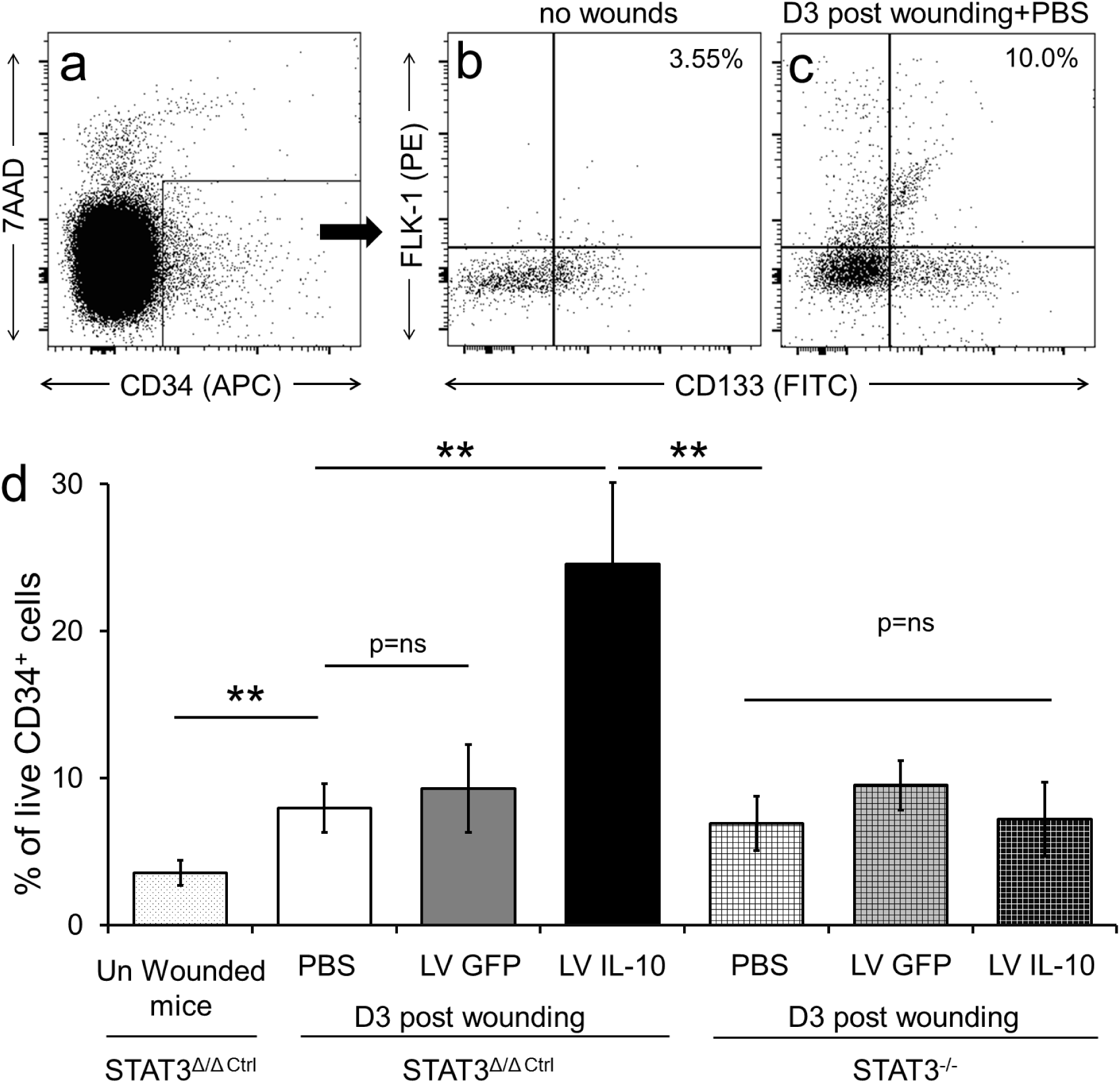
IL-10 overexpression enhanced STAT3 dependent EPC mobilization after cutaneous wounding. Peripheral blood was collected at baseline before wounding and again at day 3 post wounding from STAT3^Δ/Δ ctrl^ and STAT3^-/-^ mice that received lentiviral IL-10, lentiviral GFP or PBS treatments. Cells were stained with 7AAD, CD34, CD133 and Flk-1 for flow cytometry analysis. (a) Single cells were gated for 7AAD^-^ CD34^+^ populations; of which cells that were CD133^+^Flk-1^+^ were quantified as EPCs (b). There is a clear difference in EPC levels in peripheral blood at baseline in uninjured mice (b) vs. mice at day three post wounding (c). (d) Quantitative analysis show that following cutaneous wounding there is a significant increase in EPCs at day 3 post wounding as compared to uninjured control mice. Lentiviral IL-10 treatment significantly increase circulating EPC levels as compared to lentiviral GFP or PBS treatments in STAT3^Δ/Δ ctrl^ mice.IL-10’s effects were abrogated in STAT3^-/-^ mice. Bar plots= mean±SD of EPC numbers as analyzed by flow jo software, n=3-4 wounds from different mice/ treatment group, **p<0.01 by ANOVA.

Similar to our previous study^60^, our data shows that in comparison to unwounded mice, skin wounding and moist treatment with PBS alone resulted in a significant (p<0.01) increase in circulating levels of EPCs at day 3 post-wounding in STAT3^Δ/Δ ctrl^ mice (Figure 4b-d: 3.54±0.85 vs. 7.96±1.68; p<0.01). However, LV IL-10 overexpression in STAT3^Δ/Δ ctrl^ mice significantly increased the percentage of circulating EPC levels at day 3 post-wounding over that of mice treated with LV GFP or PBS controls (LV IL-10: 24.58±6.35 vs. LV GFP: 9.28±3.0, PBS: 7.96±1.68, p<0.01; Figure 4d). Consistent with the other observations, LV-mediated IL-10 overexpression in STAT3^-/-^ mice wounds did not experience this increase in circulating EPCs; EPC levels in these mice were analogous to LV GFP or PBS controls (STAT3^-/-^ + LV IL-10: 7.22±2.51 vs. STAT3^Δ/Δ ctrl^ + LV IL-10: 24.58±6.35, p<0.01; Figure 4d). These data show that IL-10 increased injury-induced mobilization of EPCs beyond physiologic levels, and further uphold that the function of IL- 10 overexpression in the cutaneous wound healing is STAT3-dependent.

### IL-10 increased VEGF expression via a STAT3-dependent mechanism

To investigate if IL-10-STAT3 signaling influences the expression of VEGF, a known promoter of EPC mobilization, we measured the levels of the growth factor under experimental conditions similar to those preceding. We specifically analyzed VEGF expression in the wound specimen, serum, and bone marrow (BM) as these tissue compartments represent the major microenvironments where VEGF-mediated signaling is known to facilitate injury-induced EPC mobilization. We found that, in comparison to unwounded mice, skin wounding and treatment with PBS moist treatment control significantly increased VEGF levels in all three of these compartments, most strongly in the wound itself as compared to expression levels in uninjured skin (Figure 5a-c). LV IL- 10 treatment of wounds in STAT3^Δ/Δ ctrl^ mice resulted in a further increase in VEGF levels in these three sites; while this increase trended but did not reach significance in wounded skin (Figure 5a; p=0.64), it was significant in serum (Figure 5b; p<0.01) and BM (Figure 5c; p<0.05). Once again, LV-mediated overexpression of IL-10 in STAT3^-/-^ mice did not result in increased VEGF expression.

**Figure 5:**
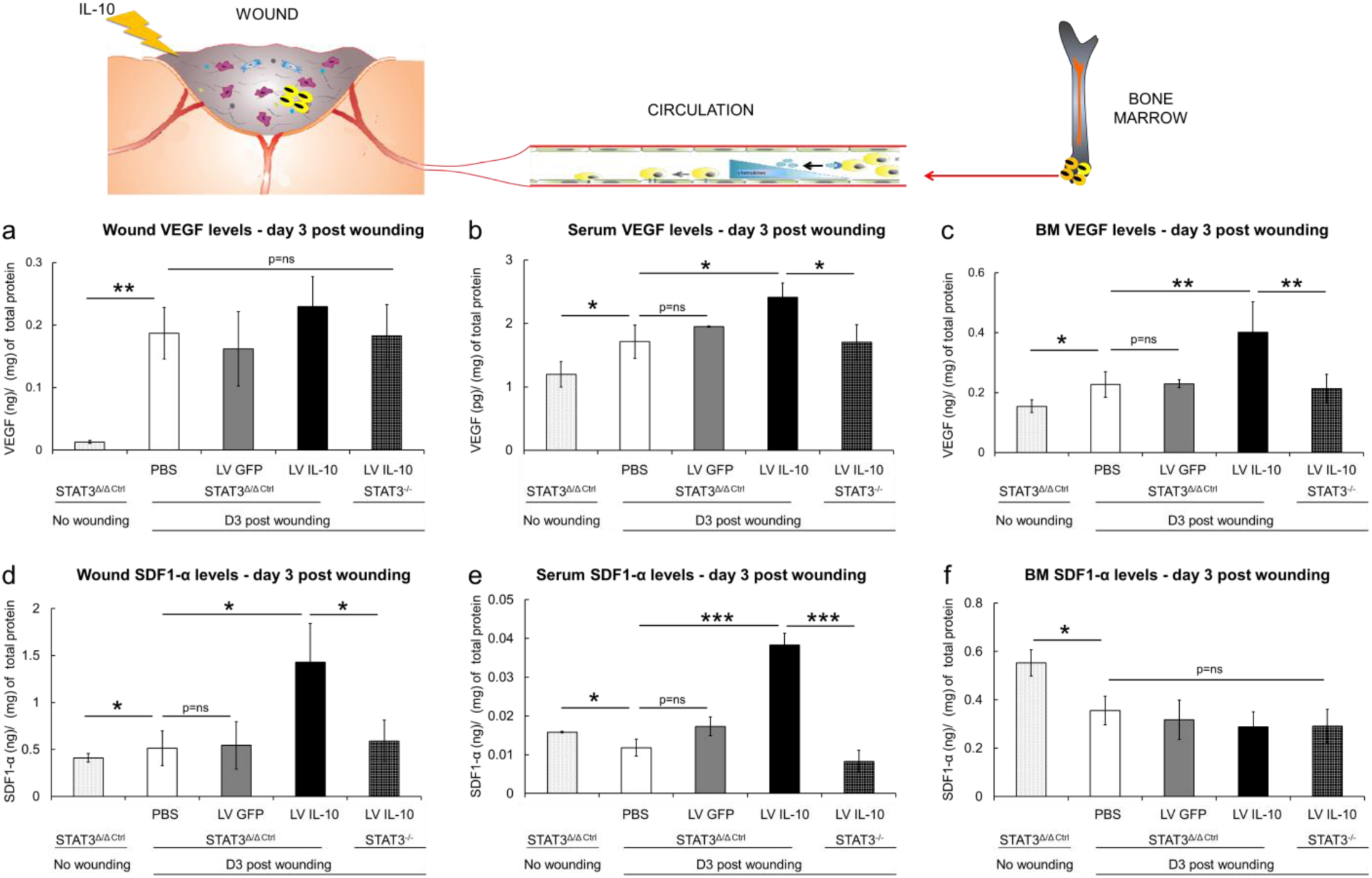
IL-10 regulates VEGF and SDF-1α gradients in wounds, serum and bone maroow in a STAT3-dependent manner. Wound tissue, peripheral blood and bone marrow (BM)were collected at baseline before wounding and again at day 3 post wounding from STAT3^Δ/Δ ctrl^ and STAT3^-/-^ mice that received lentiviral IL-10, lentiviral GFP or PBS treatments. VEGF and SDF-1α levels were quantified with ELISA. The effect of different treatments on VEGF and SDF-1α levels within the wound tissue specimens, serum and BM are respectively shown in (a-c) and (d-f). Bar plots= mean±SD, n=3-4 wounds from different animals/ treatment group, *p<0.05, **p<0.01, **p<0.001 by ANOVA.

We sought to briefly explain how LV IL-10-induced VEGF overexpression in our model might be a major contributor to increase in EPC mobilization, using a well-defined mouse model of MMP9 deficiency known to have impaired EPC mobilization after cutaneous wounding.^47^ As VEGF-induced activation of MMP9 is essential for the release of SCF and thereby EPC mobilization from the BM niche into peripheral circulation,^35^ an absence of functional MMP9 in this mouse model results in impaired EPC mobilization after cutaneous wounding. To investigate the role of VEGF/MMP9-mediated signaling in IL-10-dependent EPC mobilization, we created wounds in MMP9^-/-^ mice under similar experimental conditions as above. Consistent with previously reported findings, wounded MMP9^-/-^ mice had no change in levels of circulating EPCs at day 3 post-injury, proof of an impaired wound healing phenotype as compared to WT. As expected and as shown previously by our group, administration of SCF to MMP9^-/-^ wounded mice reinstated EPC numbers in circulation. However, when LV IL-10 was overexpressed in MMP9^-/-^ mice wounds, VEGF levels increased in the serum, but did not restore EPC mobilization (Figure S4). These data suggest that, while LV IL-10 overexpression in the wound has a systemic effect on VEGF, intact MMP9 function is likely also essential to the downstream signaling for EPC mobilization.

### IL-10 treatment resulted in a positive SDF-1α concentration gradient via a STAT3- dependent mechanism

To investigate whether the effect of IL-10 on increasing circulating EPC levels and EPC presence in dermal wounds is mediated by SDF-1α, we assessed levels of this chemotactic factor in wounds, serum, and BM at day 3 post-wounding. There was an increase in SDF-1α expression in wounded skin specimen (Figure 5d; p<0.05) at day 3 post-injury when compared to expression levels in uninjured skin. No change in serum levels of SDF-1α was seen (Figure 5e), but a significant decrease in BM SDF-1α levels (Figure 5f; p<0.01) was noted in response to cutaneous wounding. These findings support the role of SDF-1α in the retention of EPCs in a quiescent niche under homeostatic conditions, and suggest that a positive SDF-1α gradient is established following wounding, which may facilitate the egress of EPCs from the bone marrow to the circulation and homing to the site of the wound. We also observed that LV IL-10 overexpression in the wounds of STAT3^Δ/Δ ctrl^ mice at day 3 post-wounding resulted in significant increases in SDF-1α levels in wounded tissue (Figure 5d; p<0.05) and in the serum (Figure 5e; p<0.001) without affecting BM SDF-1α levels, as compared to LV GFP and PBS treatments (Figure 5f). LV IL-10’s effect in increasing SDF-1α levels in skin wounds and serum, however, was consistently abrogated in STAT3^-/-^ mice. Collectively, these data indicate that IL-10 overexpression in skin wounds establishes a STAT3- dependent SDF-1α chemotactic gradient towards the periphery, promoting EPC mobilization from the BM niche and homing to the wound site with higher local SDF-1α levels.

### IL-10 induced VEGF and SDF-1α production by fibroblasts

Because fibroblasts are important effector cells in wound healing and neovascularization, we investigated whether fibroblasts could be the source of IL-10- induced VEGF and SDF-1α expression. Consistent with this, we found that treatment with IL-10 significantly increased VEGF production in primary murine dermal fibroblasts *in vitro* (Figure 6a; p<0.05). Furthermore, the same *in vitro* treatment of primary murine dermal fibroblasts with IL-10 also resulted in significantly increased SDF-1α levels (Figure 6b; p<0.05). Together, these data provide evidence of the ability of fibroblasts to upregulate VEGF and SDF-1α levels in response to IL-10, and possibly of a central cellular role in the wound healing process observed in our *in vivo* studies.

**Figure 6:**
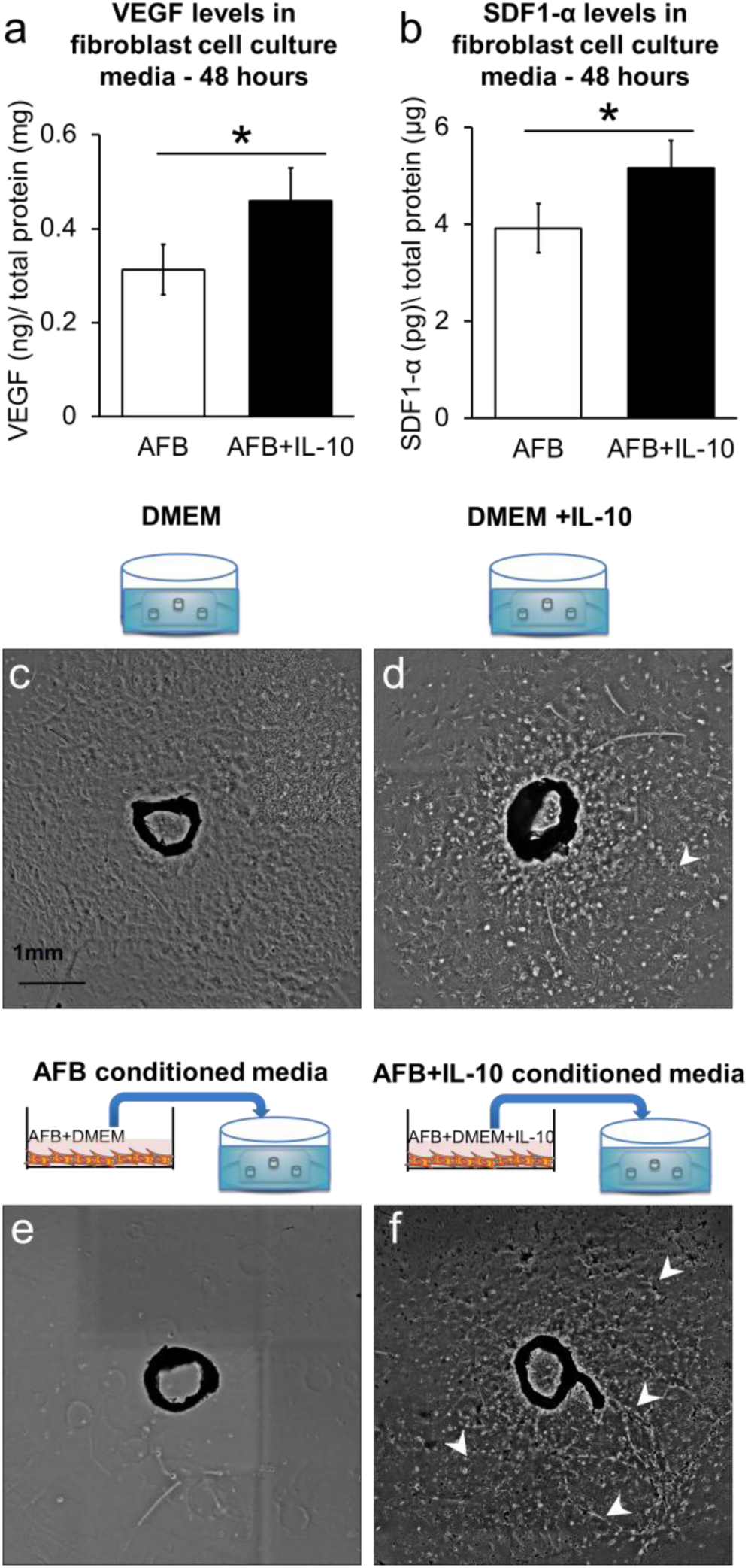
IL10 induced VEGF and SDF-1α production in primary murine dermal fibroblasts and increased cell sprouting and capillary-like network formation in aortic ring assay. Primary murine dermal fibroblasts in culture were incubated +/- 200ng/ml of IL-10 for 48 hrs. Supernatants from the cultures were assayed for VEGF and SDF-1α expression by ELISA. (a) VEGF levels significantly increase in IL 10 treated fibroblasts as compared to untreated cells. (b) SDF-1α levels significantly increase in IL- 10 treated fibroblasts as compared to controls. (c-d) Murine thoracic aortas are treated with either DMEM or DMEM supplemented with 200ng/ml IL-10 for 12 days. Aortic rings show an endothelial cell outgrowth when treated with DMEM (c), and the ones spiked with DMEM + IL10 respond similarly but with an greater relative sprouting area (d). (e-f) The aortic ring assay was repeated with either conditioned media from primary murine dermal fibroblasts, or conditioned media from fibroblasts treated with IL-10 from panel (a). The former produced no measurable endothelial outgrowth (e), whereas with the latter, a significant outgrowth with an increase in capillary-like 2D network formation was observed. White arrowheads indicate the capillary-like networks. Bar plots: mean±SD, experiments are conducted in triplicates with cells from 2 passages; *p<0.05, by ANOVA. n=3 aortic rings per treatment were studied and the experiment was repeated two times with aortas from different mice and conditioned media from different primary cell isolations.

### Relative sprouting area and lumen density on aortic ring assay is increased in IL- 10-treated fibroblast-conditioned media

The aortic ring assay is a simple but informative assay to identify modulators of angiogenesis by recapitulating the essential steps of microvessel outgrowth *in vitro*, including proliferation, migration, tubule formation, and recruitment of supporting cells. We performed this assay by comparing aortic rings cultured in DMEM complete media spiked with IL-10 versus DMEM complete media alone to determine the effect of IL-10 on capillary outgrowth. As anticipated from previous studies, aortic rings exposed to DMEM showed a robust endothelial cell (EC) outgrowth with a characteristic cobble stone-like appearance and a relative sprouting area (RSA) of 15.9±11.6mm at 12 days in culture (Figure 6c). Rings exposed to IL-10-spiked DMEM, however, had a larger RSA (24.9±12.3mm) and also showed ECs organized into 2D networks (Figure 6d; arrowheads). We then repeated the aortic ring assays with conditioned DMEM supernatant from *in vitro* cultures of untreated adult dermal fibroblasts (AFB) versus conditioned supernatant from adult dermal fibroblasts treated with IL-10. When conditioned supernatant from untreated AFB was transferred, the aortic rings showed no measurable endothelial sprouting at day 12 (Figure 6e), suggesting that factors produced by normal fibroblasts under normal cell culture conditions perhaps support vessel stabilization, as opposed to sprouting. Aortic rings exposed to conditioned supernatant from AFB cultured in DMEM+IL-10 produced sprouting with an RSA of 14.4mm±0.68mm, but interestingly, this treatment resulted in a dense capillary-like network formation (Figure 6f, arrowheads). Collectively, these data underscore the role of IL-10 in stimulating angiogenesis-inducing factors by dermal fibroblasts and suggest that fibroblasts are a potential target cell for IL-10’s pro-angiogenic effects.

### IL-10 enhanced wound healing and neovascularization in db/db murine wounds

Our data from WT mice support our working postulate that overexpression of LV IL-10 upregulates VEGF and SDF-1α, thereby promoting EPC-mediated neovascularization and wound healing. To assess the impact of this signaling under adverse wound healing conditions, we studied the effect of LV IL-10 overexpression in a db/db murine wound model of type II diabetes.

We pretreated dorsal skin in db/db mice with LV IL-10, wounding the skin after 4 days to allow transgenic protein overexpression, and then harvested these wounds at day 7 post-injury. Our results show that LV IL-10 overexpression significantly improved wound closure in db/db wounds as compared to LV GFP or PBS treated controls (Figure 7a-d). Strikingly, while the granulation tissue area was bland and comprised predominantly of adipose tissue in the LV GFP and PBS controls, LV IL-10-treated wounds had a robust granulation tissue deposition (Figure 7e), similar in morphology to WT mice wounds. We then analyzed wound neovascularization with MECA-32 staining and found a significant increase in capillary density in LV IL-10-treated db/db wounds as compared to LV GFP or PBS controls (Figure 7f-i). Furthermore, a significant increase in the CD133+Flk1+ EPC levels in the wound beds was also observed in LV IL-10-treated db/db wounds as compared to LV GFP or PBS controls (Figure 7j-m), with a concomitant increase in VEGF and SDF-1α gene expression in the wounds (Figure 7n-o). Taken together, our data support the significance of IL-10 overexpression in improving standard wound healing parameters and in increasing the recruitment, survival, and retention of EPCs to promote neovascularization and tissue repair, even under pathological conditions in which wound healing is impaired, such as in diabetes.

**Figure 7:**
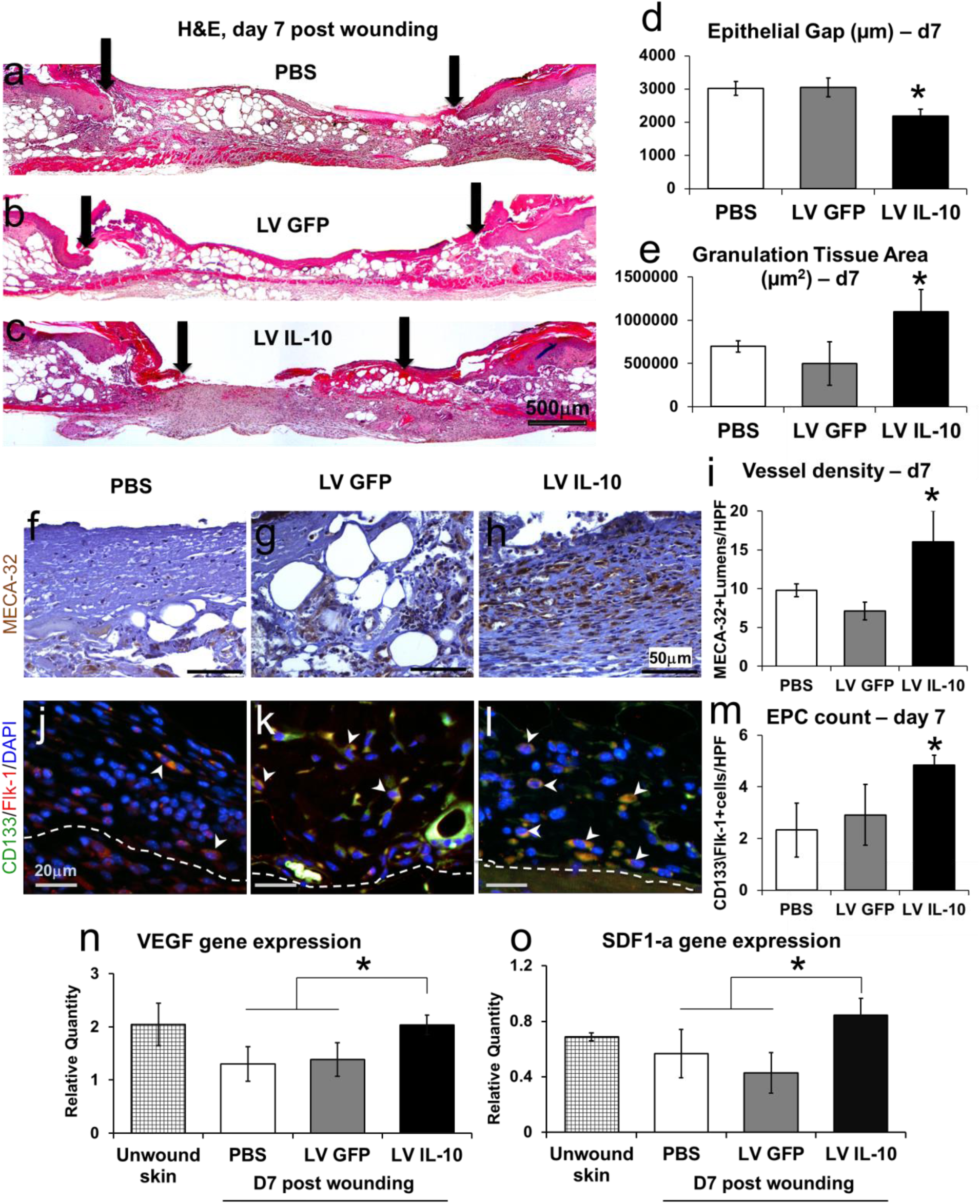
IL 10 enhanced wound healing and neovascularization in diabetic murine wound model. Dorsal wounds in db/db mice treated with either PBS, lentiviral GFP or lentiviral IL-10 were harvested at 7 days post injury. (a-c) Hematoxylin and Eosin staining reveals that lentiviral IL-10 treated wounds close faster as compared to the two control groups. Quantitation of epithelial gap (d) and granulation tissue is shown (e). (f-h) Representative images of MECA-32 stained wounds section across the three treatment conditions - PBS, lentiviral GFP or lentiviral IL-10 respectively. Increased MECA-32 stained-capillary lumens are seen in lentiviral IL-10 wounds, which is quantitatively significant (i). (j-m) Increase in the CD133+Flk1+ EPC levels in the wound beds is also observed in lentiviral IL-10-treated db/db wounds as compared to the control treatments. (n-o) qRT-PCR on wound tissue homogenates collected at day 7 shows that VEGF (n) and (o) SDF-1α expression is significantly higher IL-10 treated cohort as compared to the PBS and lentiviral GFP controls. Scale bar=500µm in (a-c), 50µm in (f-h) and 20µm in (j-l); Bar plots: mean ± SD, 2 sections/ wound, n=4 wounds from different mice/ treatment group, * p<0.05 by ANOVA.

## Discussion

In the present study, we report that IL-10 overexpression can play a pivotal role in postnatal cutaneous wound neovascularization and healing in both normal and diabetic wounds. Our data underscore a novel biological function for IL-10 in enhancing wound neovascularization by promoting EPC mobilization and recruitment, which in addition to its accepted role in immune regulation, contributes a crucial new angle to a previously reported view of regenerative cutaneous tissue repair. Moreover, our data support the capacity of IL-10 to induce EPC recruitment via VEGF and SDF-1α signaling. Therefore, we propose a model wherein IL-10 overexpression in cutaneous wounds increases the expression of VEGF and SDF-1α by fibroblasts at sites of dermal injury, resulting in a positive VEGF and SDF-1α gradient that favors mobilization of EPCs from the bone marrow, which can then home to injured tissues to support wound healing and neovascularization (Figure 8). These collective findings are also consistent with our original working hypothesis that the intracellular IL-10 signaling cascade leading to enhanced wound neovascularization and healing is STAT3-dependent^55^, as IL-10’s effects on wound neovascularization, angiogenic growth factor expression and EPC levels were lost in our murine model of skin-specific STAT3 knockdown.

**Figure 8:**
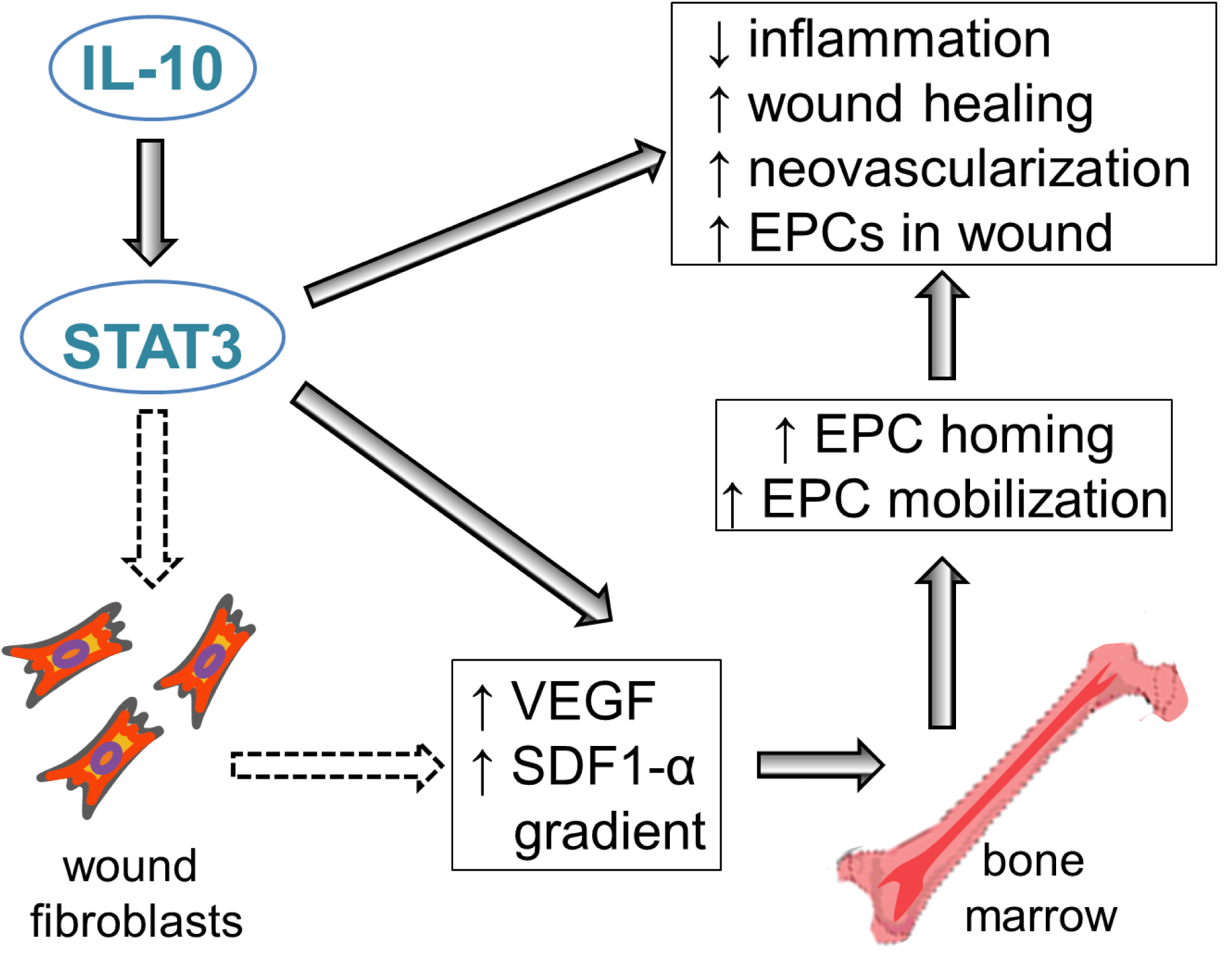
IL-10 can play a pivotal role in postnatal cutaneous wound healing and neovascularization by influencing EPC mobilization and recruitment via STAT3 dependent increase in VEGF and SDF-1α. IL10 overexpression, via a STAT3 dependent mechanism, results in enhanced levels of VEGF and SDF-1α, wound healing and neovascularization associated with an increase in EPCs in the wound. We put forth a potential pathway that IL-10 overexpression may induce wound fibroblasts to produce more VEGF and SDF-1α, which creates a positive gradient for bone marrow derived EPC mobilization and homing to healing tissue.

Importantly, our present study was also able to show the potential of IL-10 overexpression in significantly improving wound healing outcomes in diabetic mice, which included improved neovascularization and EPC recruitment and retention to the site of injury. In previous studies, the presence of unresolved inflammation in chronic wound environments has been suggested as impairing EPC function and increasing EPC apoptosis and elimination in the wound bed,^61^ a phenomenon that may partly explain why current strategies aimed at pharmacologically enhancing EPC mobilization from the BM, as well as *ex vivo* transplantation of expanded autologous EPCs^62,63^ have not attained optimal efficacy in clinical trials. In this context, IL-10 may present a more effective strategy to enhance EPC recruitment and neovascularization in impaired wound healing states such as in diabetic wounds, as it has the additional benefit of regulating wound inflammation. This strategy has been reported by Krishnamurthy *et al.*, who demonstrated in a murine myocardial infarction (MI) model that treatment with IL-10 enhanced transplanted EPC association with vascular structures and their overall survival, resulting in improved neovascularization of the injured myocardium and left ventricular function. In this study, IL-10 treatment resulted in increased STAT3 signaling and increased VEGF and SDF-1α expression in the myocardium after MI, providing mechanistic insights into IL-10’s effects on EPCs, which appear to be mediated through SDF-1α/CXCR4 and STAT3/VEGF signaling mechanisms in their model. The authors suggested that, by protecting the EPCs in an adverse microenvironment characterized by prolonged inflammation and hypoxia, IL-10 might increase the retention and numbers of these cells to participate in tissue repair,^45^ and similar mechanisms could explain the therapeutic benefit of IL-10 in promoting diabetic wound healing noted in our studies.

The present studies provide compelling evidence that treatment with IL-10 overexpression promotes VEGF expression in cutaneous wounds and establishes a positive SDF-1α concentration gradient that favors EPC mobilization from the bone marrow and homing to the site of injury, which could account for higher numbers of EPCs both in the peripheral blood and in wound microenvironment compared to baseline. Our *in vitro* data support dermal fibroblasts as one of the potential sources of IL-10-induced VEGF and SDF-1α in response to injury, something also supported by previous murine wound healing studies.^44^ However, further research is needed to fully understand the mechanism behind the IL-10-associated increases in VEGF expression observed in the serum and BM compartments. These include an assessment of the stage of molecular events, which could either place VEGF signaling upstream of SDF-1α or implicate alternative mechanisms connecting peripheral tissue injury with IL-10, VEGF, and SDF- 1α expression in the BM. In line with this rationale, our studies also propose MMP9 as an intermediary in the IL-10-induced EPC mobilization cascade, but once again, focused future studies will be required to fully assess the contribution of MMP9 to the IL-10– VEGF–SDF-1α and EPC axis. Evidence of a direct link between BM-derived MMP9 and VEGF-induced angiogenesis, including BM cell mobilization and BM-derived cell recruitment to angiogenic foci in the brain has been demonstrated,^64^ but whether IL-10 is influential in facilitating this angiogenic response in cutaneous wounding is yet to be determined. Nevertheless, the reciprocal relationship between IL-10, MMP-9, and VEGF warrants further investigation in light of the findings in this study. These studies could provide relevant information that could eventually lead to the design of innovative targeted therapies to optimize EPC mobilization, wound neovascularization, and regenerative wound healing. Cell types other than fibroblasts could also be involved in the effects reported here, and future efforts will be directed towards adapting our skin-specific STAT3 knockout model system to develop keratinocyte, fibroblast, or endothelial cell specific STAT3-deficient mice. The findings from these models can then be compared to the skin-specific knockdown to better understand the contribution by the individual skin compartments.

A role for IL-10 in dermal wound repair has been established in the literature. Sato et al demonstrated that IL-10 expression peeked about 3 hours after wounding and once again at 3 days after injury, with keratinocytes and mononuclear cells being the major sources of IL-10^65^. Skin wound healing in a fetal skin transplant model from *IL-10^-/-^* mice suggested that IL-10 is essential for fetal scarless tissue repair^66^. These studies suggest an important role for endogenous IL-10 in regulating inflammatory cell infiltration and cytokine production in cutaneous wound healing. Given the notion that while IL-10 expression rapidly increases in adult wounds at 3 hours post wounding, and yet postnatal cutaneous wounds heal with scar formation as compared to their mid-gestation fetal counterpart, it is plausible that relevant physiological levels cannot be upheld under adult dermal conditions. Indeed, IL-10 is an immunoregulatory cytokine that limits both innate as well as adaptive immune responses of the host-immune interactions, mainly aimed to protect the host from immune-mediated tissue damage. IL-10 is known to inhibit overexpression of broad spectrum of pro-inflammatory cytokines and chemokines such as IL-1, IL-6, IL-8, TNF-α, monocyte chemoattractant protein MCP-1, macrophage inflammatory protein (MIP)-1 in wound healing^65,66^. We and others have shown that IL-10 overexpression regulates inflammatory responses to promote a regenerative wound healing phenotype in murine cutaneous wounds in a dose-dependent manner.^51–53^ Furthermore, we have recently reported a novel role for IL-10 in inducing a fibroblast- mediated formation of an extracellular wound matrix rich in hyaluronan^54^, which proved to be essential for IL-10’s regenerative wound healing capability. Several other studies have also shown that IL-10 regulates the synthesis and degradation of various extracellular matrix molecules in different fibroblastic cell types, which upholds the notion of potential IL-10 anti-fibrotic mechanisms in the skin^67–69^. Yet, its impact on the process of cutaneous wound neovascularization in physiologic and pathophysiologic disease states such as diabetes has not been completely deciphered. A study by Eming *et al.* ^70^ reported that in comparison to WT controls, cutaneous wounds of IL-10^-/-^ mice show an accelerated angiogenic response during the early phase (day 3) of wound healing, along with an increase in VEGF-A. However, by day 5, the angiogenic rush and VEGF-A upregulation subside with no difference in vessel density in healed wounds. In this study, macrophages comprise the majority of VEGF-A-expressing cells in IL-10^-/-^ wounds, and though overall healing was rapid, these wounds were characterized by poor strength, increased macrophage infiltration, increased angiogenic burst, increase in α-sma smooth muscle cell (SMA) positive myofibroblast-mediated wound contraction, and increased wound collagen/scar tissue. These observations may arguably be interpreted as compensatory mechanisms initiated by the lack of anti-inflammatory IL-10 in these knockout mice. Similar findings of an increase in angiogenic burst along with robust inflammation and rapid closure of wounds were reported in mutant mice deficient in the expression of other chemokines, such as IL-12/23^71^. Furthermore, IL-10–deficient EPCs had impaired migratory function *in vitro* in response to SDF-1α. Such impairments in local SDF-1α expression levels and EPC function have also been noted in diabetic wound healing. ^44^ Further inquiry must be undertaken to elucidate whether IL10 directly alters the biology of EPCs leading to their greater survival and function with roles in physiologic and constitutively inflamed diseased states.

In summary, our data provide evidence that supports the role for IL-10 in enhancing VEGF and SDF-1α levels, neovascularization, EPC recruitment, and ultimately improving cutaneous wound healing outcomes via STAT3 signaling. The immunoregulatory and anti-inflammatory effects of IL-10 may also serve to enhance EPC mobilization, survival, and function in the wounds. These data additionally point out to the need to design precision strategies to deliver targeted and sustained levels of IL-10 with the therapeutic potential to enhance EPC-driven angiogenesis and wound healing in normal and diabetic wounds.

## List of abbreviations

ΔΔCt: Comparative Ct
4-OHT: 4-hydroxy tamoxifen
AFB: Adult Fibroblasts
ANOVA: Analysis of Variance
BGS: Bovine Growth Serum
BM: Bone marrow
CD34: cluster of differentiation 34 molecule
CD133: cluster of differentiation 133 molecule (also known as Prominin1)
CXCR4: CXC chemokine receptor type 4
DMEM: Dulbecco’s Modified Eagle’s Media
ECL: Enhanced Chemoluminescence
ECs: Endothelial cells
ECM: Extracellular Matrix
ELISA: Enzyme-Linked Immunosorbent Assay
EPCs: Endothelial progenitor cells
FLK-1: Fetal Liver Kinase 1 (also known as KDR, is a receptor for VEGF)
GFP: Green Fluorescent Protein
H&E: Hematoxylin and Eosin
HIF-1: Hypoxia inducible factor 1
HSCs: Hematopoietic stem cells
IgG: Immunoglobulin G
IHC: Immunohistochemistry
IL-10: Interleukin 10
IL-10R: Interleukin 10 Receptor
LV-GFP: Lenti Virus expressing GFP
LV-IL-10: Lenti Virus expressing IL-10
MI: Myocardial Infarction
PBS: Phosphate Buffered Saline
PSF: Penicillin, Streptomycin, Amphotericin
p-STAT3: Phosphorylated Signal Transducer and Activator of Transcription 3
qPCR: Real Time-Polymerase Chain Reaction
SDF-1α: Stromal cell-derived factor 1α
shRNA: Short Hairpin RNA
STAT3: Signal Transducer and Activator of Transcription 3
TBS: Tris-Buffered Saline\
TGF-β1, β3: Transforming Growth Factor-β1, β3
Tween-20: Polysorbate-20
VEGF: Vascular endothelial growth factor

## Conflict of Interest

The authors on the manuscript do not have any conflict of interest (either financial or personal).

## Author contributions

S.B., E.S., T.M.C., P.B.L., and S.G.K. designed the experiments; S.B., E.S., X.W., H.V.V., N.T, A.B., C.M.M., performed bench and animal related experimental work; S.B., E.S., X.W., H.V.V., N.T, A.B., C.M.M., D.A.N., M.J.B., T.M.C., P.B.L., and S.G.K. analyzed data; S.B., E.S., X.W., H.V.V., N.T, A.B., C.M.M., D.A.N., T.M.C., M.J.B., P.B.L., and S.G.K. contributed to the discussions, manuscript writing and editing. We affirm that all authors have read and agree with the manuscript.

## Acknowledgements and Funding Sources

The authors sincerely acknowledge the technical support received from our laboratory staff members and the support received from Cincinnati Children’s Hospital Medical Center Flow Cytometry Core. The authors also acknowledge the editorial support of Drs. Monica Fahrenholtz and Hector Martinez-Valdez from the Office of Surgical Research Administration (OSRA), Department of Surgery, Texas Children’s Hospital. This study is supported by 1R01GM111808-01 NIH/NIGMS (SGK) and Wound Healing Society Foundation 3M Award (SB, SGK).

## Notes

The authors declare no competing financial interests.

## References

1. Economic Costs of Diabetes in the U.S. in 2017. Diabetes care. May 2018;41(5):917–928.

2. Frykberg RG, Banks J. Challenges in the Treatment of Chronic Wounds. Advances in wound care. Sep 1 2015;4(9):560–582.

3. Casqueiro J, Alves C. Infections in patients with diabetes mellitus: A review of pathogenesis. Indian journal of endocrinology and metabolism. Mar 2012;16 Suppl 1:S27–36.

4. Balaji S, King A, Crombleholme TM, Keswani SG. The Role of Endothelial Progenitor Cells in Postnatal Vasculogenesis: Implications for Therapeutic Neovascularization and Wound Healing. Advances in wound care. Jul 2013;2(6):283–295.

5. Ribatti D, Vacca A, Nico B, Roncali L, Dammacco F. Postnatal vasculogenesis. Mechanisms of development. Feb 2001;100(2):157–163.

6. Asahara T, Masuda H, Takahashi T, et al. Bone marrow origin of endothelial progenitor cells responsible for postnatal vasculogenesis in physiological and pathological neovascularization. Circulation research. Aug 6 1999;85(3):221–228.

7. Urbich C, Dimmeler S. Endothelial progenitor cells: characterization and role in vascular biology. Circulation research. Aug 20 2004;95(4):343–353.

8. Tepper OM, Capla JM, Galiano RD, et al. Adult vasculogenesis occurs through in situ recruitment, proliferation, and tubulization of circulating bone marrow-derived cells. Blood. Feb 1 2005;105(3):1068–1077.

9. Carmeliet P. Developmental biology. One cell, two fates. Nature. Nov 2 2000;408(6808):43, 45.

10. Ehrbar M, Metters A, Zammaretti P, Hubbell JA, Zisch AH. Endothelial cell proliferation and progenitor maturation by fibrin-bound VEGF variants with differential susceptibilities to local cellular activity. Journal of controlled release: official journal of the Controlled Release Society. Jan 3 2005;101(1-3):93–109.

11. Urbich C, Aicher A, Heeschen C, et al. Soluble factors released by endothelial progenitor cells promote migration of endothelial cells and cardiac resident progenitor cells. Journal of molecular and cellular cardiology. Nov 2005;39(5):733–742.

12. Patel J, Seppanen EJ, Rodero MP, et al. Functional Definition of Progenitors Versus Mature Endothelial Cells Reveals Key SoxF-Dependent Differentiation Process. Circulation. Feb 21 2017;135(8):786–805.

13. Asahara T, Murohara T, Sullivan A, et al. Isolation of putative progenitor endothelial cells for angiogenesis. Science. Feb 14 1997;275(5302):964–967.

14. Peichev M, Naiyer AJ, Pereira D, et al. Expression of VEGFR-2 and AC133 by circulating human CD34(+) cells identifies a population of functional endothelial precursors. Blood. Feb 1 2000;95(3):952–958.

15. Falanga V. Wound healing and its impairment in the diabetic foot. Lancet. Nov 12 2005;366(9498):1736–1743.

16. Blakytny R, Jude E. The molecular biology of chronic wounds and delayed healing in diabetes. Diabet Med. Jun 2006;23(6):594–608.

17. Phillips TJ. Chronic cutaneous ulcers: etiology and epidemiology. The Journal of investigative dermatology. Jun 1994;102(6):38S–41S.

18. Kim KA, Shin YJ, Kim JH, et al. Dysfunction of endothelial progenitor cells under diabetic conditions and its underlying mechanisms. Archives of pharmacal research. Feb 2012;35(2):223–234.

19. Capla JM, Grogan RH, Callaghan MJ, et al. Diabetes impairs endothelial progenitor cell-mediated blood vessel formation in response to hypoxia. Plastic and reconstructive surgery. Jan 2007;119(1):59–70.

20. Madonna R, De Caterina R. Cellular and molecular mechanisms of vascular injury in diabetes--part II: cellular mechanisms and therapeutic targets. Vascular pharmacology. Mar-Jun 2011;54(3-6):75–79.

21. Callaghan MJ, Ceradini DJ, Gurtner GC. Hyperglycemia-induced reactive oxygen species and impaired endothelial progenitor cell function. Antioxidants & redox signaling. Nov-Dec 2005;7(11-12):1476–1482.

22. Fadini GP, Miorin M, Facco M, et al. Circulating endothelial progenitor cells are reduced in peripheral vascular complications of type 2 diabetes mellitus. Journal of the American College of Cardiology. May 3 2005;45(9):1449–1457.

23. Keswani SG, Katz AB, Lim FY, et al. Adenoviral mediated gene transfer of PDGF- B enhances wound healing in type I and type II diabetic wounds. Wound Repair Regen. Sep-Oct 2004;12(5):497–504.

24. Tepper OM, Galiano RD, Capla JM, et al. Human endothelial progenitor cells from type II diabetics exhibit impaired proliferation, adhesion, and incorporation into vascular structures. Circulation. Nov 26 2002;106(22):2781–2786.

25. Vasa M, Fichtlscherer S, Aicher A, et al. Number and migratory activity of circulating endothelial progenitor cells inversely correlate with risk factors for coronary artery disease. Circulation research. Jul 6 2001;89(1):E1–7.

26. Kalka C, Masuda H, Takahashi T, et al. Transplantation of ex vivo expanded endothelial progenitor cells for therapeutic neovascularization. Proceedings of the National Academy of Sciences of the United States of America. Mar 28 2000;97(7):3422–3427.

27. Suh W, Kim KL, Kim JM, et al. Transplantation of endothelial progenitor cells accelerates dermal wound healing with increased recruitment of monocytes/macrophages and neovascularization. Stem Cells. Nov-Dec 2005;23(10):1571–1578.

28. Balaji S, Vaikunth SS, Lang SA, et al. Tissue-engineered provisional matrix as a novel approach to enhance diabetic wound healing. Wound repair and regeneration: official publication of the Wound Healing Society [and] the European Tissue Repair Society. Jan-Feb 2012;20(1):15–27.

29. Cho H, Balaji S, Sheikh AQ, et al. Regulation of endothelial cell activation and angiogenesis by injectable peptide nanofibers. Acta biomaterialia. Jan 2012;8(1):154–164.

30. Georgescu A, Alexandru N, Constantinescu A, Titorencu I, Popov D. The promise of EPC-based therapies on vascular dysfunction in diabetes. European journal of pharmacology. Nov 1 2011;669(1-3):1–6.

31. Li B, Sharpe EE, Maupin AB, et al. VEGF and PlGF promote adult vasculogenesis by enhancing EPC recruitment and vessel formation at the site of tumor neovascularization. FASEB journal: official publication of the Federation of American Societies for Experimental Biology. Jul 2006;20(9):1495–1497.

32. Asahara T, Takahashi T, Masuda H, et al. VEGF contributes to postnatal neovascularization by mobilizing bone marrow-derived endothelial progenitor cells. The EMBO journal. Jul 15 1999;18(14):3964–3972.

33. Rosti V, Massa M, Campanelli R, De Amici M, Piccolo G, Perfetti V. Vascular endothelial growth factor promoted endothelial progenitor cell mobilization into the peripheral blood of a patient with POEMS syndrome. Haematologica. Sep 2007;92(9):1291–1292.

34. Heeschen C, Dimmeler S, Hamm CW, Boersma E, Zeiher AM, Simoons ML. Prognostic significance of angiogenic growth factor serum levels in patients with acute coronary syndromes. Circulation. Feb 4 2003;107(4):524–530.

35. Heissig B, Hattori K, Dias S, et al. Recruitment of stem and progenitor cells from the bone marrow niche requires MMP-9 mediated release of kit-ligand. Cell. May 31 2002;109(5):625–637.

36. Tang JM, Wang JN, Zhang L, et al. VEGF/SDF-1 promotes cardiac stem cell mobilization and myocardial repair in the infarcted heart. Cardiovascular research. Aug 1 2011;91(3):402–411.

37. Sengupta N, Afzal A, Caballero S, et al. Paracrine modulation of CXCR4 by IGF-1 and VEGF: implications for choroidal neovascularization. Investigative ophthalmology & visual science. May 2010;51(5):2697–2704.

38. Peled A, Petit I, Kollet O, et al. Dependence of human stem cell engraftment and repopulation of NOD/SCID mice on CXCR4. Science. Feb 5 1999;283(5403):845–848.

39. Rey M, Valenzuela-Fernandez A, Urzainqui A, et al. Myosin IIA is involved in the endocytosis of CXCR4 induced by SDF-1alpha. Journal of cell science. Mar 15 2007;120(Pt 6):1126–1133.

40. Liu ZJ, Tian R, An W, et al. Identification of E-selectin as a novel target for the regulation of postnatal neovascularization: implications for diabetic wound healing. Annals of surgery. Oct 2010;252(4):625–634.

41. Moore MA, Hattori K, Heissig B, et al. Mobilization of endothelial and hematopoietic stem and progenitor cells by adenovector-mediated elevation of serum levels of SDF-1, VEGF, and angiopoietin-1. Annals of the New York Academy of Sciences. Jun 2001;938:36–45; discussion 45-37.

42. Hattori K, Heissig B, Tashiro K, et al. Plasma elevation of stromal cell-derived factor-1 induces mobilization of mature and immature hematopoietic progenitor and stem cells. Blood. Jun 1 2001;97(11):3354–3360.

43. Bauer SM, Bauer RJ, Liu ZJ, Chen H, Goldstein L, Velazquez OC. Vascular endothelial growth factor-C promotes vasculogenesis, angiogenesis, and collagen constriction in three-dimensional collagen gels. Journal of vascular surgery. Apr 2005;41(4):699–707.

44. Gallagher KA, Liu ZJ, Xiao M, et al. Diabetic impairments in NO-mediated endothelial progenitor cell mobilization and homing are reversed by hyperoxia and SDF-1 alpha. The Journal of clinical investigation. May 2007;117(5):1249–1259.

45. Krishnamurthy P, Thal M, Verma S, et al. Interleukin-10 deficiency impairs bone marrow-derived endothelial progenitor cell survival and function in ischemic myocardium. Circulation research. Nov 11 2011;109(11):1280–1289.

46. Losordo DW, Dimmeler S. Therapeutic angiogenesis and vasculogenesis for ischemic disease. Part I: angiogenic cytokines. Circulation. Jun 1 2004;109(21):2487–2491.

47. Cho H, Balaji S, Hone NL, et al. Diabetic wound healing in a MMP9-/- mouse model. Wound Repair Regen. Sep 2016;24(5):829–840.

48. Couper KN, Blount DG, Riley EM. IL-10: the master regulator of immunity to infection. J Immunol. May 1 2008;180(9):5771–5777.

49. Ouyang W, Rutz S, Crellin NK, Valdez PA, Hymowitz SG. Regulation and functions of the IL-10 family of cytokines in inflammation and disease. Annual review of immunology. 2011;29:71–109.

50. Wang Y, Fan L, Meng X, et al. Transplantation of IL-10-transfected endothelial progenitor cells improves retinal vascular repair via suppressing inflammation in diabetic rats. Graefe’s archive for clinical and experimental ophthalmology = Albrecht von Graefes Archiv fur klinische und experimentelle Ophthalmologie. Oct 2016;254(10):1957–1965.

51. Peranteau WH, Zhang L, Muvarak N, et al. IL-10 overexpression decreases inflammatory mediators and promotes regenerative healing in an adult model of scar formation. The Journal of investigative dermatology. Jul 2008;128(7):1852–1860.

52. Gordon A, Kozin ED, Keswani SG, et al. Permissive environment in postnatal wounds induced by adenoviral-mediated overexpression of the anti-inflammatory cytokine interleukin-10 prevents scar formation. Wound Repair Regen. Jan-Feb 2008;16(1):70–79.

53. Kieran I, Knock A, Bush J, et al. Interleukin-10 reduces scar formation in both animal and human cutaneous wounds: results of two preclinical and phase II randomized control studies. Wound Repair Regen. May-Jun 2013;21(3):428–436.

54. Balaji S, Wang X, King A, et al. Interleukin-10-mediated regenerative postnatal tissue repair is dependent on regulation of hyaluronan metabolism via fibroblast-specific STAT3 signaling. FASEB journal: official publication of the Federation of American Societies for Experimental Biology. Mar 2017;31(3):868–881.

55. Moore KW, de Waal Malefyt R, Coffman RL, O’Garra A. Interleukin-10 and the interleukin-10 receptor. Annual review of immunology. 2001;19:683–765.

56. Levy DE, Lee CK. What does Stat3 do? The Journal of clinical investigation. May 2002;109(9):1143–1148.

57. Hiramatsu K, Sasagawa S, Outani H, Nakagawa K, Yoshikawa H, Tsumaki N. Generation of hyaline cartilaginous tissue from mouse adult dermal fibroblast culture by defined factors. The Journal of clinical investigation. Feb 2011;121(2):640–657.

58. Baker M, Robinson SD, Lechertier T, et al. Use of the mouse aortic ring assay to study angiogenesis. Nature protocols. Dec 22 2011;7(1):89–104.

59. Pfaffl MW. A new mathematical model for relative quantification in real-time RT-PCR. Nucleic Acids Res. May 1 2001;29(9):e45.

60. Morris LM, Klanke CA, Lang SA, et al. Characterization of endothelial progenitor cells mobilization following cutaneous wounding. Wound Repair Regen. Jul-Aug 2010;18(4):383–390.

61. Desouza CV, Hamel FG, Bidasee K, O’Connell K. Role of inflammation and insulin resistance in endothelial progenitor cell dysfunction. Diabetes. Apr 2011;60(4):1286–1294.

62. Ingram DA, Caplice NM, Yoder MC. Unresolved questions, changing definitions, and novel paradigms for defining endothelial progenitor cells. Blood. Sep 1 2005;106(5):1525–1531.

63. Fadini GP, Losordo D, Dimmeler S. Critical reevaluation of endothelial progenitor cell phenotypes for therapeutic and diagnostic use. Circulation research. Feb 17 2012;110(4):624–637.

64. Hao Q, Su H, Palmer D, et al. Bone marrow-derived cells contribute to vascular endothelial growth factor-induced angiogenesis in the adult mouse brain by supplying matrix metalloproteinase-9. Stroke. Feb 2011;42(2):453–458.

65. Sato Y, Ohshima T, Kondo T. Regulatory role of endogenous interleukin-10 in cutaneous inflammatory response of murine wound healing. Biochemical and biophysical research communications. Nov 1999;265(1):194–199.

66. Liechty KW, Kim HB, Adzick NS, Crombleholme TM. Fetal wound repair results in scar formation in interleukin-10-deficient mice in a syngeneic murine model of scarless fetal wound repair. Journal of pediatric surgery. Jun 2000;35(6):866–872; discussion 872-863.

67. Yamamoto T, Eckes B, Krieg T. Effect of interleukin-10 on the gene expression of type I collagen, fibronectin, and decorin in human skin fibroblasts: differential regulation by transforming growth factor-beta and monocyte chemoattractant protein-1. Biochemical and biophysical research communications. Feb 16 2001;281(1):200–205.

68. Moroguchi A, Ishimura K, Okano K, Wakabayashi H, Maeba T, Maeta H. Interleukin-10 suppresses proliferation and remodeling of extracellular matrix of cultured human skin fibroblasts. European surgical research. Europaische chirurgische Forschung. Recherches chirurgicales europeennes. Jan-Feb 2004;36(1):39–44.

69. Shi JH, Guan H, Shi S, et al. Protection against TGF-beta1-induced fibrosis effects of IL-10 on dermal fibroblasts and its potential therapeutics for the reduction of skin scarring. Archives of dermatological research. May 2013;305(4):341–352.

70. Eming SA, Werner S, Bugnon P, et al. Accelerated wound closure in mice deficient for interleukin-10. The American journal of pathology. Jan 2007;170(1):188–202.

71. Matias MA, Saunus JM, Ivanovski S, Walsh LJ, Farah CS. Accelerated wound healing phenotype in Interleukin 12/23 deficient mice. J Inflamm (Lond). Dec 20 2011;8:39.

