## supplemental figures and legends for "IL-10 Promotes Endothelial Progenitor Cell Driven Wound Neovascularization and Enhances Healing via STAT3"

**Supplemental Figure Legends**

**Figure S1: Validation of the skin-specific conditional STAT3 Knockout murine model.** Dorsal skin of STAT-3$^{\Delta/\Delta}$ mice was topically treated with 4-OHT or vegetable oil (vehicle control) for 7 days and harvested. Western Blot (a) and densitometric analysis (b) demonstrate ~60% knockdown of STAT-3 with 4-OHT application. Bar plots: mean±SD, n=4 different mice/ treatment group, *** $p<0.001$ by ANOVA.

**Figure S2: Increased IL-10 expression and stat3 phosphorylation in lentiviral IL-10 treated wounds.** (a) wound tissue homogenates from day 3 post wounding show a significant increase in IL-10 expression levels in lentiviral IL-10 treated mice as compared to PBS and lentiviral GFP controls with ELISA. (b-c) Representative images from immunohistochemical staining analysis of wounds in STAT-3$^{\Delta/\Delta\ ctrl}$ transgenic mice treated with lentiviral IL-10 show STAT-3 and phospho-STAT-3 signaling. Bar plots: mean±SD, n=4 wounds from different mice/ treatment group, * $p<0.05$ by ANOVA.

**Figure S3: IL-10 overexpression enhanced wound remodeling and collagen deposition in a STAT3 dependent manner.** Movat's pentachrome staining of wounds at day 7 post wounding in STAT-3$^{\Delta/\Delta}$ mice. Original magnification: 4X (a-d); (a-c) similar granulation tissue was observed between the three groups – PBS, lentiviral GFP and lentiviral IL-10 treated wounds; however IL-10 treated cohort (c) displayed greater collagen deposition in the center of the wound and increased epidermal thickness as compared to the control groups (a-b) in STAT-3$^{\Delta/\Delta\ ctrl}$ mice. The effect of IL-10 overexpression on granulation tissue collagen is not apparent in STAT-3$^{-/-}$ mice.

**Figure S4: MMP9 signaling is necessary for IL-10-dependent EPC mobilization. Dorsal wounds were created in MMP9$^{-/-}$ mice.** Peripheral blood was collected at day 3 post wounding from WT and MMP9$^{-/-}$ mice that received lentiviral IL-10, lentiviral GFP or PBS treatments. Cells were stained with 7AAD, CD34, CD133 and Flk-1 for flow cytometry analysis. (a) Single cells were gated for 7AAD$^{-}$ CD34$^{+}$ populations; of which cells that were CD133$^{+}$Flk-1$^{+}$ were quantified as EPCs. Wounded MMP9$^{-/-}$ mice had no change in levels of circulating EPCs at day 3 post-injury, proof of an impaired wound healing phenotype as compared to background strain controlled WT mice. As expected and as shown previously by our group, administration of SCF to MMP9$^{-/-}$ wounded mice reinstated EPC numbers in circulation. However, lentiviral IL-10 overexpressed in MMP9$^{-/-}$ mice wounds did not increase EPC mobilization. (b) Analysis of the serum levels of VEGF with different treatments. Bar plots: mean±SD, n=4 mice/ treatment group, * $p<0.01$ by ANOVA.

**Supplemental Figures**

**Supplemental Figure 1**

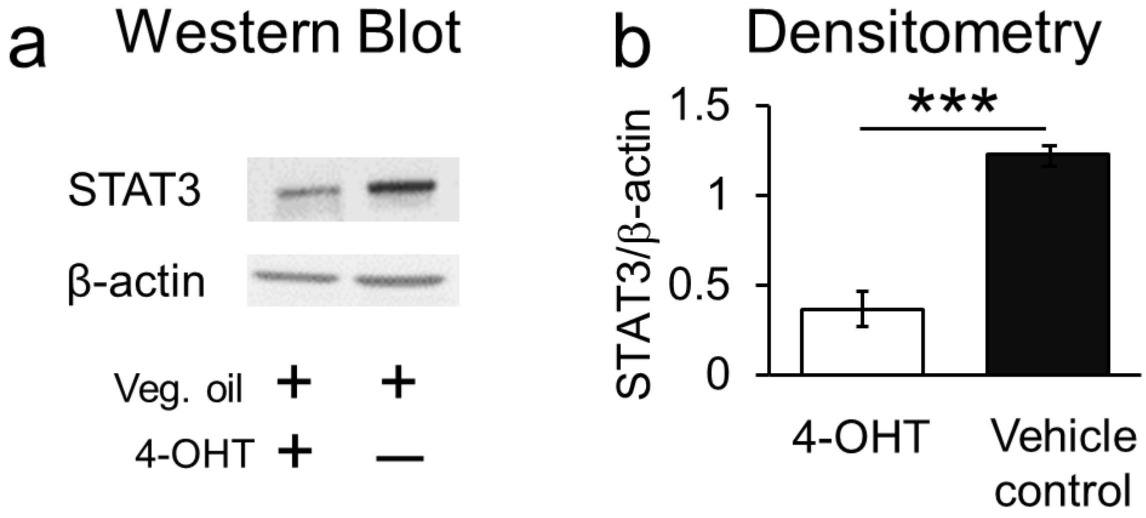

**Supplemental Figure S2**

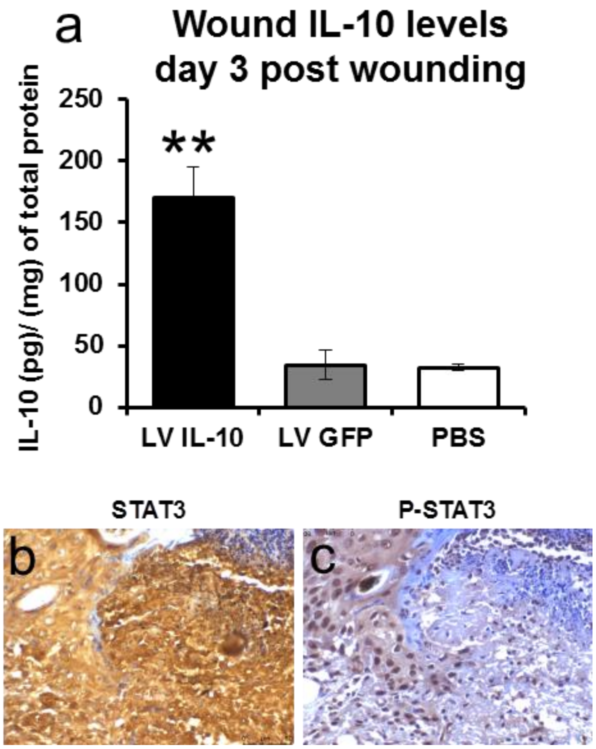

**Supplemental Figure 3**

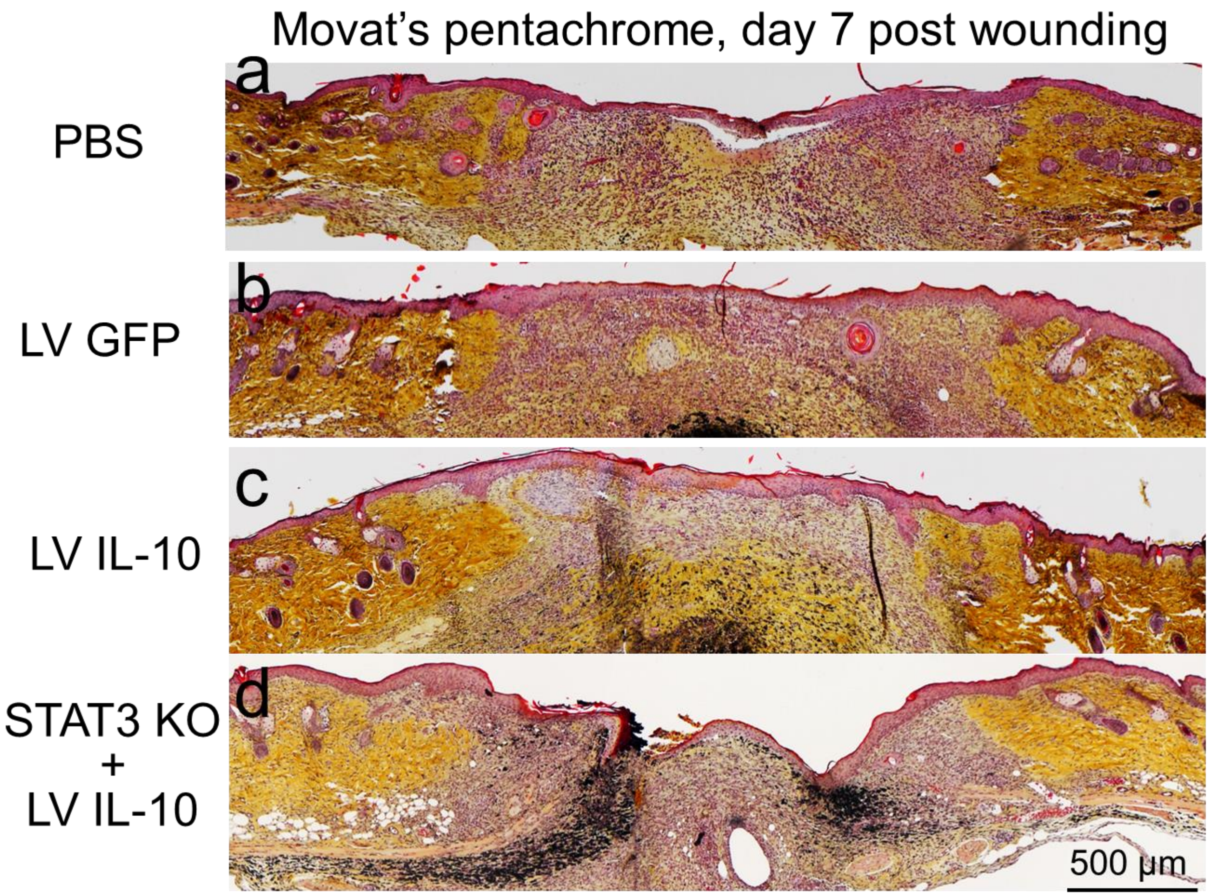

### Supplemental Figure 4

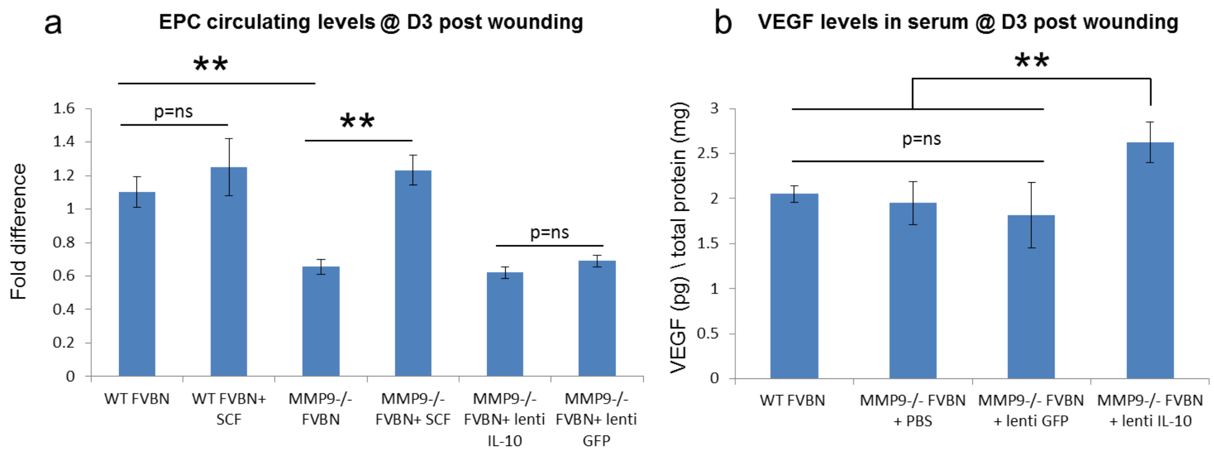
